# Serotonergic neurons in the dorsal raphe nucleus scale O₂ utilization in response to traumatic stress

**DOI:** 10.64898/2026.01.14.699586

**Authors:** Yuan Zhang, Yi-da Pan, Wen-ying Zheng, Qin-qing Wen, Min-zhen Zhu, Huan-yu Li, Wen-quan Liang, Zheng-yan Zhu, Bao-chen Liu, Xin-rui Yuan, Yu Wu, Jing-hua Dong, Maria Antonietta Davoli, Gustavo Turecki, Naguib Mechawar, Xin-hong Zhu

## Abstract

Trauma is a main cause of psychiatric disorders. Approximately 24% of trauma-exposed individuals suffer from posttraumatic stress disorder throughout their lifetime, however, the vast majority of people are resilient. Why some people are resilient while others develop mental disorders in face of trauma in a short timescale? Here, we show that O₂ actively initiates a cell-signaling pathway that regulates behavioral outcomes in response to traumatic stress. We found that serotonergic neurons in the dorsal raphe nucleus (DRN^5-HT^ neurons) projecting to the ventral caudate putamen and the ventral tegmental area exhibit distinct physiological patterns of O₂ utilization. In response to traumatic stress, O₂ utilization in these neurons changed individually, following either a susceptible or resilient pattern, thereby modulating corresponding behaviors. This O₂ scaling was regulated by ferroptosis within the DRN^5-HT^ neurons. Thus, we identify a previously unrecognized function of O_2_, opening an avenue for the treatment and prevention of stress-related disorders.

## INTRODUCTION

Traumatic stress has been implicated in the development of multiple psychiatric disorders, including posttraumatic stress disorder (PTSD) and major depressive disorder (MDD) ^1,2^. Although approximately 24% of trauma-exposed individuals experience PTSD at some point in their lifetime ^3^, most people remain resilient—maintaining mental health or recovering rapidly after temporary impairments ^4,5^. Both depression and resilience are adaptive behavioral responses that can persist over time; however, affective experiences—defined as an individual’s emotional reaction to an event—are brief and occur rapidly ^5^. Thus, trauma quickly triggers behavioral changes, but how it produces distinct, even opposite, behavioral outcomes within such a short timescale remains unclear.

Oxygen (O₂), which readily diffuses across cell membranes and the blood–brain barrier, may play a role in emotional regulation. Since its discovery in 1774, O₂ has been recognized as a fundamental gas essential for all living organisms. Cells must therefore detect and respond to O₂ deficiency to survive this life-threatening condition. The first identified and best-characterized oxygen-sensing mechanism is the <u>h</u>ypoxia-inducible factor 1α (HIF1α)-dependent <u>o</u>xygen-<u>s</u>ensing <u>p</u>athway (HOSP) in metazoans ^6–10^. HIF1α accumulates under hypoxic conditions and directly activates transcription of more than 300 genes that enable cellular adaptation to low-oxygen environments. Under normoxic conditions, HIF1α proteins are hydroxylated by dioxygenases—primarily prolyl hydroxylase 2 (PHD2) and factor-inhibiting HIF1 (FIH1)—and subsequently recognized by ubiquitin ligase complexes, typically von Hippel–Lindau (VHL), for degradation.

Evidence suggests that the HOSP contributes to the pathophysiology of PTSD and MDD ^11–14^. Our previous studies demonstrated that intermittent hypobaric hypoxia training increases the expression of HIF1α target genes in the dorsal raphe nucleus (DRN) and enhances resilience in adult mice ^15–17^. In addition, NADPH oxidase 4 (NOX4) is the predominant oxygen-consuming enzyme in the brain and constitutively generates hydrogen peroxide (H₂O₂)—a reactive oxygen species (ROS) that physiologically inhibits PHD2/FIH1 activity at nanomolar concentrations—thereby modulating HOSP activation ^18,19^. A genome-wide association study identified a region downstream of *NOX4* associated with multiple neuropsychiatric disorders, including PTSD and MDD ^20^. Furthermore, NOX4 expression can be regulated by sirtuin 1, encoded by the depression-associated gene *SIRT1* ^21–23^.

At sea level, the partial pressure of oxygen (*p*O₂) in the Earth’s atmosphere is approximately 160 mmHg (∼21% of air), whereas within the brain, *p*O₂ ranges from 1–40 mmHg ^24,25^. Although oxygen-sensing mechanisms in living organisms are well established ^6–10^, the physiological implications of such low and heterogeneous *p*O₂ in the brain remain largely unknown. Here, we investigated whether and how O₂ in the brain regulates emotional processing. We demonstrate that O₂ actively initiates a cell-signaling pathway that influences behavioral outcomes in response to traumatic stress. Specifically, we found that serotonergic neurons in the DRN (hereafter referred to as DRN^5-HT^ neurons) projecting to the ventral caudate putamen (vCPU) and the ventral tegmental area (VTA) differentially regulate depression-related behaviors through the NOX4–HOSP axis. The basal levels of NOX4-dependent O₂ consumption in these DRN^5-HT^ neurons vary under physiological conditions. Following traumatic stress, O₂ utilization in these neurons shifts toward either a susceptible (SUS) or resilient (RES) pattern, thereby modulating corresponding behavioral outcomes. This scaling of O₂ utilization is regulated by ferroptosis within the DRN^5-HT^ neurons. Our findings provide insight into oxygen-mediated signaling in the brain and suggest new avenues for treating and preventing stress-related disorders.

## RESULTS

### O₂ utilization in DRN^5-HT^ neurons is altered in depression

Directly measuring changes in cellular O₂ utilization in the brain is challenging. NOX4 consumes O₂ to generate physiological levels of H₂O₂, which subsequently activates the HOSP pathway (Fig. 1a). Previous studies have shown that dihydroethidium (DHE) is the most suitable probe for detecting ROS in mouse brains with low *p*O₂ levels ^26,27^ (Supplementary Fig. 1a). To determine whether cellular O₂ utilization contributes to emotional regulation, we performed whole-brain imaging using sagittal brain slices to assess DHE fluorescence intensity in mice subjected to a 10-day chronic social defeat stress (CSDS) paradigm—a well-established mouse model of PTSD. In this paradigm, male C57BL/6J mice are repeatedly exposed to larger, highly aggressive resident CD1 mice allowing classification into SUS or RES groups ^28,29^. We observed that DHE fluorescence intensity was lower in many brain regions of RES mice compared with controls, except in the DRN, where DHE intensity specifically decreased in SUS mice but remained unchanged in RES mice. Coronal brain slices confirmed this selective decrease in the DRN of SUS mice (Supplementary Fig. 1b–1h). Combined with our previous findings that the DRN is highly sensitive to hypoxia ^16,17^, these results suggest that cellular O₂ utilization in the DRN contributes to behavioral outcomes following traumatic stress.

**Fig. 1:**
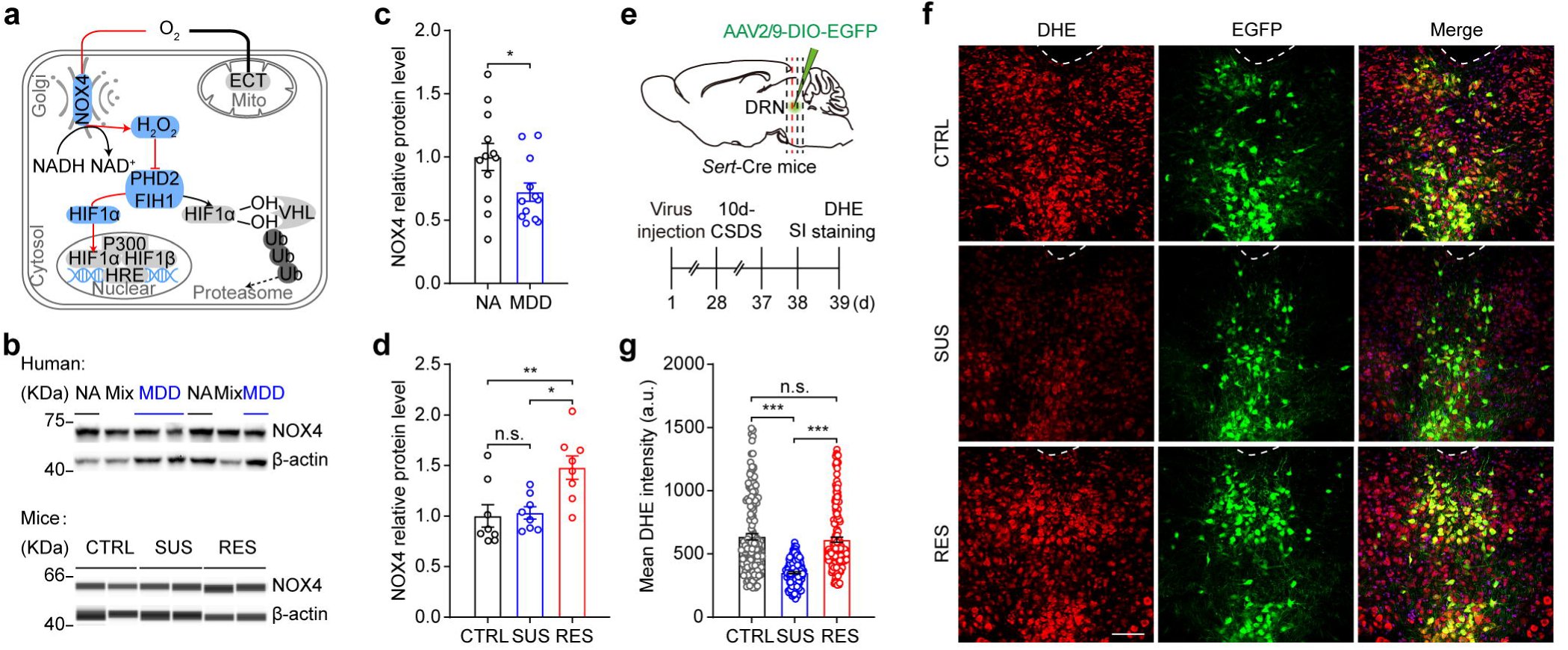
Cellular O₂ utilization is altered in the DRN of patients with MDD and mice following CSDS. **a.** Schematic representation of the NOX4–HOSP axis. Oxygen is used by NOX4 to produce H₂O₂, which stabilizes HIF1α by inhibiting the HIF hydroxylases, PHD2/FIH1. **b.** Representative western blot bands for NOX4 and β-actin. The tissue level of β-actin was used as the internal standard, and a sample of mixed human brain tissues (Mix) was used as a loading control. Protein expression was normalized to the levels of the internal standard and mixture. MDD, individuals with major depressive disorder; NA, matched accidental- or natural-death controls; SUS, susceptible mice following a 10-day CSDS paradigm; RES, resilient mice; CTRL, control mice not subjected to CSDS. **c.** Quantification of NOX4 in the DRN of individuals with MDD and matched controls (*n* = 12). **d.** Protein levels of NOX4 in the DRN of SUS, RES, and CTRL mice following a 10-day CSDS paradigm (*n* = 8). **e.** Schematic of virus injection into the DRN of *Sert*-Cre mice. The dotted red lines represent the bregma of the brain slices shown in (**f**). **f.** Oxidized DHE fluorescence in DRN^5-HT^ neurons after a 10-day CSDS paradigm. **g.** Quantification of cellular DHE fluorescence intensity in DRN^5-HT^ neurons (*n* = 6 mice per phenotype; each dot represents the DHE fluorescence intensity in one cell). Scale bars, 100 μm; \**p* < 0.05, \*\**p* < 0.01, and \*\*\**p* < 0.001. **c,** two-tailed unpaired *t*-test. **d, g,** one-way analyses of variance (ANOVA) with Tukey’s multiple comparisons test.. n.s., not significant. Data are presented as mean ± SEM. Statistical analysis parameters are included in Supplemental Table 4.

NOX4 is the principal oxygen-consuming enzyme in the brain that constitutively produces H₂O₂ at nanomolar concentrations ^30^. We next examined NOX4 expression in the DRN of postmortem human brain tissue from individuals with MDD. NOX4 protein levels were significantly lower in the DRN of subjects with MDD than in nondepressed controls (Fig. 1b and 1c). Notably, after the 10-day CSDS paradigm, NOX4 protein levels were unchanged in the DRN of SUS mice but significantly upregulated in RES mice compared with controls (Fig. 1b and 1d). These data indicate that alterations in NOX4-driven O₂ utilization in the DRN may underlie behavioral responses to traumatic stress.

The DRN is the major source of serotonergic neurons in the brain ^31^. To explore the specific role of O₂ utilization by DRN^5-HT^ neurons in depression, we injected an AAV2/9-DIO-EGFP virus into the DRN of *Sert*-Cre mice—a mouse line in which Cre recombinase activity is restricted to DRN^5-HT^ neurons ^32^—and subjected the mice to the 10-day CSDS paradigm (Fig. 1e). DHE staining revealed that DHE fluorescence intensity was significantly reduced in EGFP⁺ DRN^5-HT^ neurons of SUS mice, whereas no significant change was observed in those of RES mice compared with controls (Fig. 1f and 1g). This pattern of DHE fluorescence in *Sert*-Cre mice mirrored that observed in C57BL/6J mice following CSDS (Supplementary Fig. 1g and 1h). In contrast, the DHE fluorescence intensity of GABAergic neurons in the DRN (DRNᴳᴬᴮᴬ) was decreased in both SUS and RES mice relative to controls (Supplementary Fig. 2a–2d). These findings suggest that altered O₂ consumption by DRN^5-HT^ neurons, rather than DRNᴳᴬᴮᴬ neurons, plays a dominant role in mediating behavioral responses to CSDS.

**Fig. 2:**
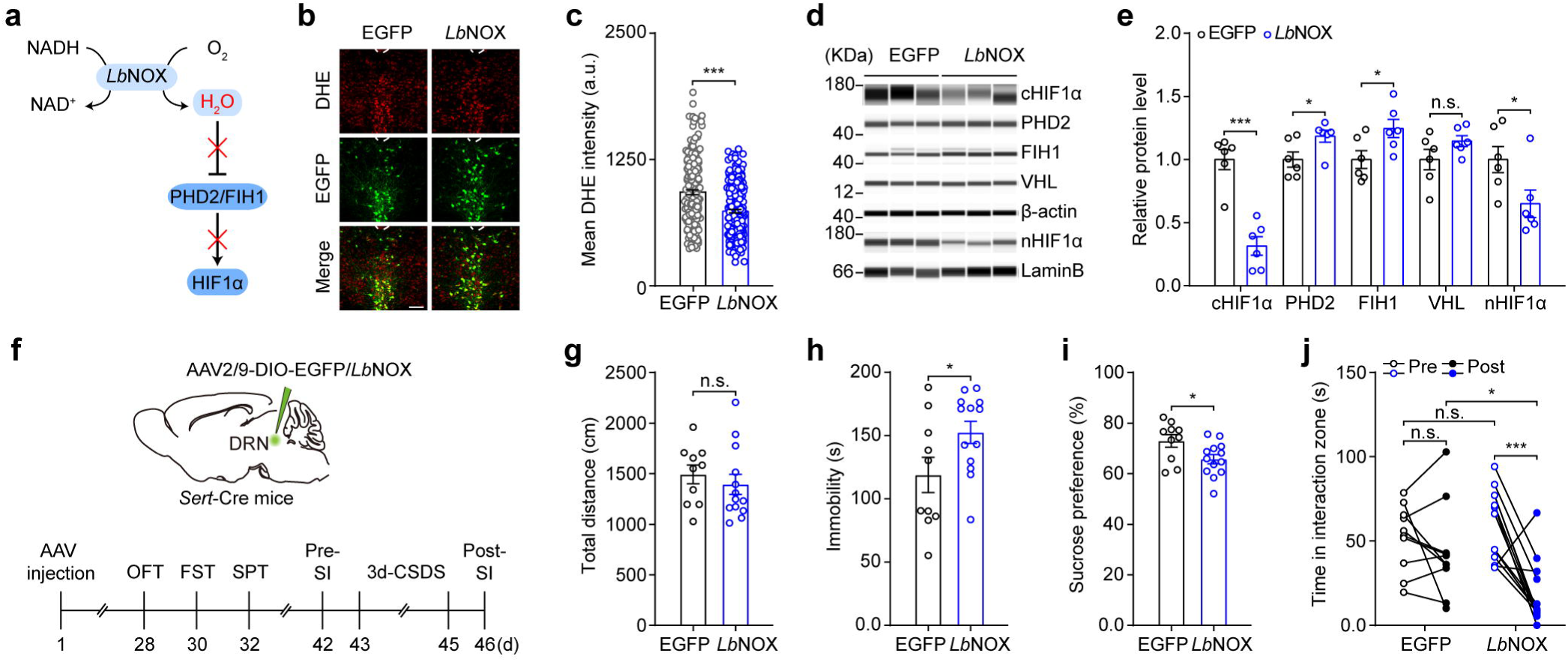
*Lb*NOX expression in DRN^5-HT^ neurons induces depressive-like behaviors. **a.** Working model of *Lb*NOX. *Lb*NOX couples the oxidation of NADH to NAD⁺ with the reduction of O₂ to water and thus disrupts the NOX4–HOSP axis. **b, c.** Oxidized DHE staining revealed lower ROS levels in the DRN of mice with *Lb*NOX expression than in *Sert*-Cre mice injected with AAV2/9-DIO-EGFP (EGFP) into the DRN (*n* = 6 mice per group). **d, e.** Representative immunoblot images and quantification of the protein levels related to the HOSP normalized to loading controls (*n* = 6). **f–j.** Experimental timeline for injections of AAV2/9-DIO-*Lb*NOX and control virus into the DRN of *Sert*-Cre mice and behavioral tests (*n*_EGFP_ = 10, *n_Lb_*_NOX_ = 13). Total distance traveled in the open field test (OFT) (**g**), immobility in the forced swimming test (FST) (**h**), ratio of sucrose preference in the sucrose preference test (SPT) (**i**), and social interaction time before (Pre) or after (Post) the 3-day CSDS paradigm (**j**). Scale bars, 100 μm; \**p* < 0.05, \*\**p* < 0.01, and \*\*\**p* < 0.001. **c, g, h, i,** two-tailed unpaired *t*-test. **e,** multiple *t*-test. **j,** two-way ANOVA with Sidak’s multiple comparisons test. n.s., not significant. Data are presented as mean ± SEM.

To further characterize the role of cellular O₂ utilization in DRN^5-HT^ neurons in behavioral regulation, we modulated O₂ consumption genetically using *Lactobacillus brevis* NADH oxidase (*Lb*NOX)—a water-forming bacterial oxidase that couples NADH oxidation to NAD⁺ regeneration and reduces O₂ to water ^33,34^. This reaction increases cellular O₂ utilization while disrupting the NOX4–HOSP signaling pathway (Fig. 2a). To validate this manipulation, we microinjected AAV2/9-DIO-*Lb*NOX or a control virus into the DRN of *Sert*-Cre mice and sacrificed the animals 4 weeks later (Supplementary Fig. 2e). Expression of *Lb*NOX in DRN^5-HT^ neurons significantly decreased the cellular NADH/NAD⁺ ratio in the DRN compared with controls (Supplementary Fig. 2f), consistent with previous findings ^33,34^. Notably, cellular DHE fluorescence intensity in DRN^5-HT^ neurons expressing *Lb*NOX was lower than that in controls (Fig. 2b and 2c). Furthermore, *Lb*NOX expression markedly increased protein levels of PHD2 and FIH1 while reducing both cytosolic and nuclear levels of HIF1α in the DRN (Fig. 2d, 2e and Supplementary Fig. 2g). These findings indicate that *Lb*NOX expression elevates O₂ consumption in DRN^5-HT^ neurons while suppressing activation of the HOSP.

After validating this approach, we conducted behavioral assessments in a new cohort of *Sert*-Cre mice 4 weeks after viral injection (Fig. 2f). *Lb*NOX expression in DRN^5-HT^ neurons significantly increased immobility time in the forced swim test (FST) without affecting locomotor activity in the open field test (OFT) relative to controls (Fig. 2g and 2h). In addition, *Lb*NOX expression significantly reduced sucrose preference in the sucrose preference test (SPT) (Fig. 2i). During a 3-day subthreshold CSDS paradigm, *Sert*-Cre mice expressing *Lb*NOX exhibited social avoidance—spending less time in the interaction zone—whereas control mice behaved similarly to unstressed controls (Fig. 2j). These findings demonstrate that *Lb*NOX expression in DRN^5-HT^ neurons induces depressive-like behaviors. In a parallel experiment, we found that *Lb*NOX expression in DRNᴳᴬᴮᴬ neurons had no measurable effects on behavioral responses (Supplementary Fig. 2h–2l). Together, these results indicate that O₂-driven cellular signaling in DRN^5-HT^ neurons specifically regulates behavioral responses to traumatic stress.

### The NOX4–HOSP axis in DRN^5-HT^ neurons is essential for depression

Our previous findings indicated that HIF1α in the DRN plays an important role in the pathophysiology of depression ^15–17^. To further investigate the contribution of the HOSP to depressive behavior, we first examined the expression of HOSP-related genes in the DRN of mice following a 10-day CSDS paradigm. Compared with controls, *Hif1a* mRNA levels were significantly increased, whereas *Hif1an* (encoding FIH1) mRNA levels were decreased in the DRN of RES mice. Similarly, nuclear HIF1α protein levels were significantly elevated, and FIH1 expression was downregulated in the DRN of RES mice relative to controls. No significant changes were detected in the DRN of SUS mice. In a separate model of PTSD—the stress-enhanced fear learning (SEFL) paradigm ^35^—we observed activation of the HOSP in the DRN of RES but not SUS mice (Fig. 3a–3c, Supplementary Fig. 3a and 3b).

**Fig. 3:**
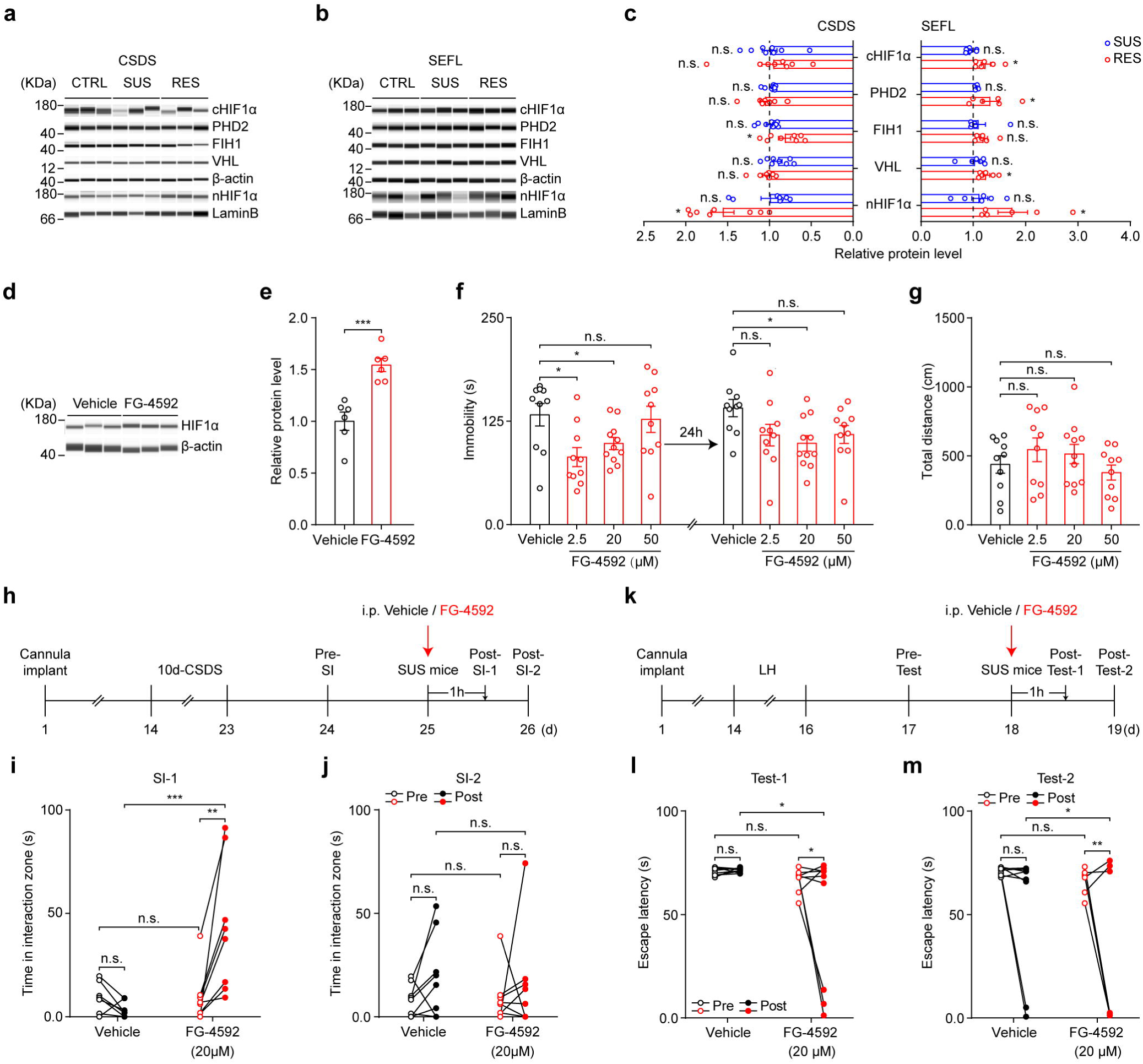
Infusion of FG-4592 into the DRN produces antidepressant-like effects in multiple animal models of depression. **a–c.** Western blot images (**a** and **b**) and quantification (**c**) of protein levels of HIF1α, PHD2, and FIH1 in the DRN of adult C57BL/6J mice subjected to a 10-day CSDS (*n* = 8) or SEFL paradigm (*n* = 6). cHIF1α, cytosolic HIF1α; nHIF1α, nuclear HIF1α. Lamin B, a marker of the nucleus. **d, e.** Western blot and quantification of HIF1α and β-actin levels in the DRN following DRN-infusion of FG-4592 (20 μM) (*n* = 6). **f.** Immobility time in the FST after infusion of FG-4592 at different concentrations or vehicle into the DRN of C57BL/6J mice 1 h or 24 h after treatment (*n* = 10–11). **g.** Total distance travelled in the OFT 24 h after FG-4592 treatment (*n* = 10–11). **h–j.** Experimental timeline for DRN-infusion of FG-4592 (20 μM) in CSDS. After cannula implantation, C57BL/6J mice were subjected to a 10-day CSDS paradigm, and SUS mice were given only one injection of FG-4592, and behaviors were detected 1 h or 24 h after treatment. Time in the social interaction zone 1 h (SI-1) (**i**) and 24 h (SI-2) (**j**) after the infusion of FG-4592 (*n* = 8). **k–m.** Experimental procedure for DRN infusion of FG-4592 (20 μM) in the learning helplessness (LH) paradigm. Following the LH paradigm, SUS mice were used to evaluate the effects of DRN infusion of FG-4592 (single injection) on behavioral changes. Escape latency was evaluated 1 h (Test-1) (**l**) or 24 h (Test-2) (**m**) after FG-4592 treatment (*n* = 9). FG-4592 application significantly decreased the latency to escape in the LH paradigm compared to control. \**p* < 0.05, \*\**p* < 0.01, and \*\*\**p* < 0.001. **c, g,** one-way ANOVA with Tukey’s multiple comparisons test. **e,** two tailed unpaired *t*-test. **f,** two-way repeated measures (RM) ANOVA with Sidak’s multiple-comparisons test. **i, j, l, m,** two-way ANOVA with Sidak’s multiple-comparisons test. n.s., not significant. Data are presented as mean ± SEM.

To investigate whether the HOSP is a useful target for drug screening, we infused FG-4592 (roxadustat), a clinically used inhibitor of HIF prolyl hydroxylase ^36^, into the DRNs of C57BL/6J mice, and found that a single-dose infusion of FG-4592 (20 μM) significantly increased the protein levels of HIF1α in the DRN with minimal neuronal toxicity (Fig. 3d, 3e and Supplementary Fig. 3c). Then, we infused a single dose of FG-4592 at different concentrations or vehicle into the DRN of adult C57BL/6J mice and conducted FST at 1 and 24 h after infusion. We found that infusion of FG-4592 at concentrations of 2.5 and 20 μM, respectively, significantly shortened immobility 1 h after the infusion. Notably, the antidepressant-like effect of FG-4592 infusion at the concentration of 20 μM was lasting 24 h, and DRN-infusion of FG-4592 had little effect on locomotion (Fig. 3f and 3g). Moreover, when using the CSDS paradigm, including FG-4592 or vehicle infusion into the DRN of SUS mice, we found that FG-4592 reversed the social avoidance induced by CSDS paradigm (Fig. 3h–3j and Supplementary Fig. 3d–3f). Similarly, DRN-infusion of FG-4592 (20 μM) produced a rapid and long-lasting antidepressant-like effects following learning helplessness paradigms (Fig. 3k–3m). These results are consistent with our previous reports ^15,16^, indicating that HOSP activation in the DRN is involved in the pathophysiology of depression.

Next, to determine the functional importance of HIF1α, we microinjected AAV2/9-DIO-EGFP-*Hif1a*-shRNA into the DRN of *Sert-Cre* mice to downregulate HIF1α in a Cre-dependent manner ^16^. Expression of *Hif1a-*shRNA in DRN^5-HT^ neurons induced a depressive-like phenotype, characterized by increased immobility in the FST without changes in locomotion, reduced sucrose preference, and social avoidance following a 3-day CSDS paradigm (Fig. 4a–4e). In contrast, infusion of AAV2/9-DIO-EGFP-*Hif1an*-shRNA or AAV2/9-DIO-EGFP-*Egln1*-shRNA into the DRN—each of which upregulates HIF1α expression—produced antidepressant-like effects in both the FST and 10-day CSDS paradigms (Supplementary Fig. 4a–4j). Notably, the depressive phenotypes induced by *Lb*NOX expression in DRN^5-HT^ neurons were completely reversed by simultaneous downregulation of FIH1 and PHD2 in these neurons (Fig. 4f–4m). In parallel, downregulation of HIF1α in DRNᴳᴬᴮᴬ neurons had minimal effects on depression-related behaviors (Supplementary Fig. 4k–4o). Together, these results clearly demonstrate that the NOX4–HOSP axis in DRN^5-HT^ neurons is essential for the development of depressive behavior.

**Fig. 4:**
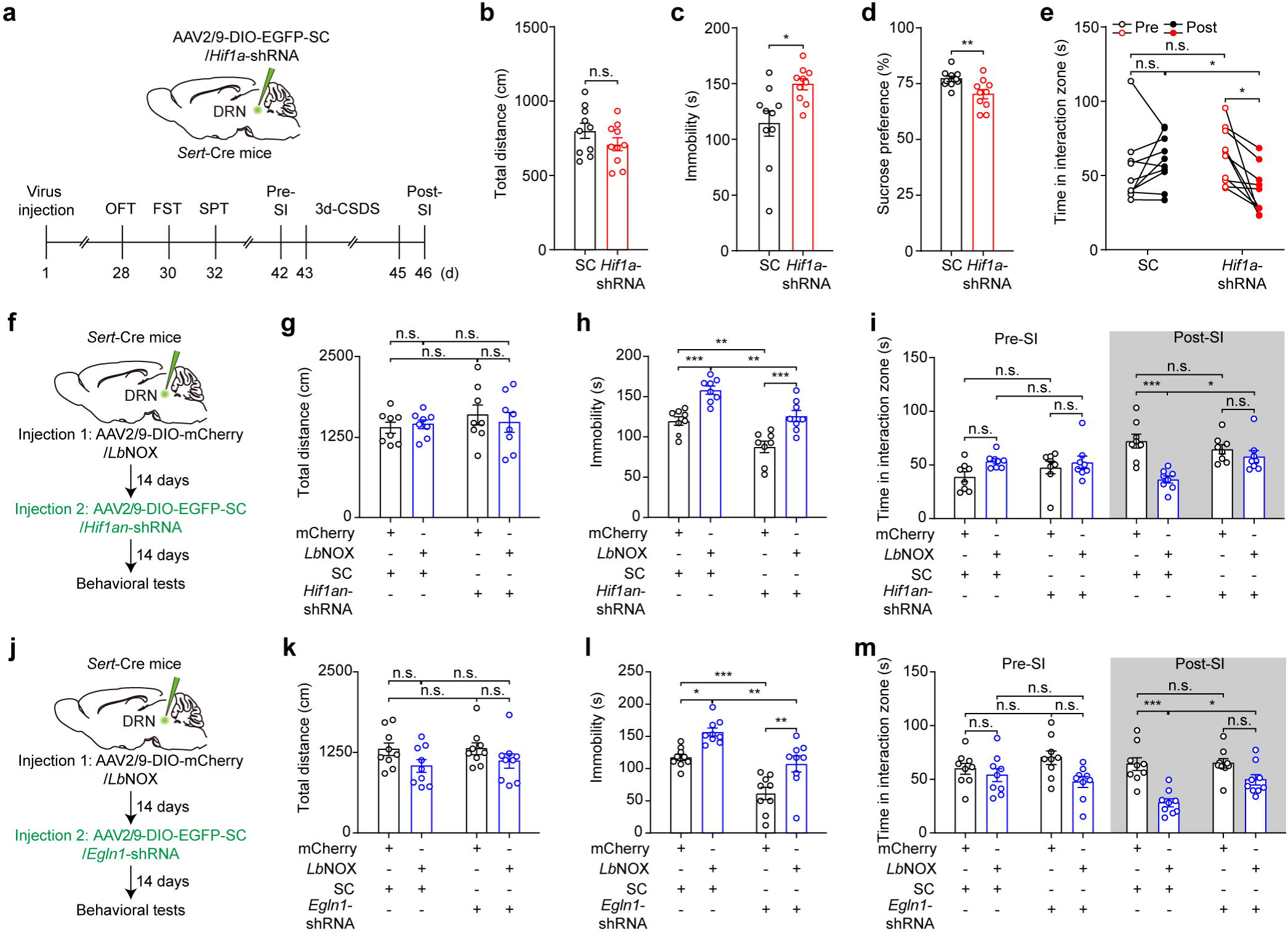
The NOX4–HOSP axis in DRN^5-HT^ neurons is essential for depression. **a–e.** Behavioral tests following *Hif1a* knockdown in DRN^5-HT^ neurons (*n* = 10). Experimental timeline (**a**). Total distance traveled in the OFT (**b**), immobility in the FST (**c**), SPT ratio (**d**), and social interaction time before (Pre) or after (Post) a 3-day CSDS paradigm (**e**). **f–m.** Experimental timelines for rescue experiments in which FIH1 (*n* = 8) (**f–i**) or PHD2 (*n* = 9) (**j–m**) was knocked down in DRN^5-HT^ neurons from *Sert*-Cre mice injected with *Lb*NOX or control virus (mCherry). Total distance traveled in the OFT (**g** and **k**), immobility in the FST (**h** and **l**), and social interaction time measured before and after a 3-day CSDS paradigm (**i** and **m**). SC, control virus with scrambled sequence. \**p* < 0.05, \*\**p* < 0.01, and \*\*\**p* < 0.001. **b, c, d,** two-tailed unpaired *t*-test. **e,** two-way ANOVA with Sidak’s multiple comparisons. **g, h, i, k, l, m,** two-way ANOVA with Tukey’s multiple comparisons. n.s., not significant. Data are presented as mean ± SEM.

### DRN^5-HT^ neurons differentially regulate depression-related behaviors via the HOSP

To investigate the neural circuit mechanisms underlying O₂ regulation in the DRN in response to traumatic stress, we performed a whole-brain analysis to evaluate the concentrations of 5-HT and its metabolites in *Sert*-Cre mice injected with AAV2/9-DIO-EGFP-*Hif1a*-shRNA or a control virus in the DRN. We found that expression of *Hif1a*-shRNA in DRN^5-HT^ neurons significantly decreased 5-HT concentrations in many brain areas, including the medial prefrontal cortex (mPFC), vCPU, VTA, and medial geniculate body (MGN), compared with those in control animals, with little effect on its metabolites. To determine which brain area receiving serotonergic input mediates depression-related behavior via the HOSP, we microinjected AAV2/9-DIO-EGFP-*Hif1a*-shRNA into the DRN of C57BL/6J mice and a retrograde AAV2/R-Cre virus into the aforementioned brain regions. We found that expression of *Hif1a*-shRNA in DRN neurons projecting to the VTA (DRN→VTA neurons) specifically decreased immobility in the FST without affecting locomotion in the OFT, indicating that downregulation of HIF1α in DRN→VTA neurons produced an antidepressant-like effect. This hypothesis was verified using a 10-day CSDS paradigm and the novelty-suppressed feeding test (NSFT), a behavioral assay used to assess the efficacy of chronic antidepressant treatment ^15,37^, in which downregulation of HIF1α in DRN→VTA neurons significantly decreased the latency to approach food without affecting food consumption compared with that of control animals. In addition, although DRN infusion of AAV2/9-DIO-EGFP-*Hif1a*-shRNA had little effect on immobility in the FST in mice injected with AAV2/R-Cre into the vCPU, possibly due to increased locomotor activity in the OFT, downregulation of HIF1α in DRN→vCPU neurons significantly decreased the sucrose preference ratio in experimental mice compared with control mice. Moreover, these mice displayed depression-like behaviors in the NSFT and a 3-day CSDS paradigm, indicating that downregulation of HIF1α in DRN→vCPU neurons induced a depressive phenotype (Supplementary Fig. 5a–5k). These results indicate that DRN^5-HT^ neurons differentially regulate depression-related behaviors via the HOSP.

**Fig. 5:**
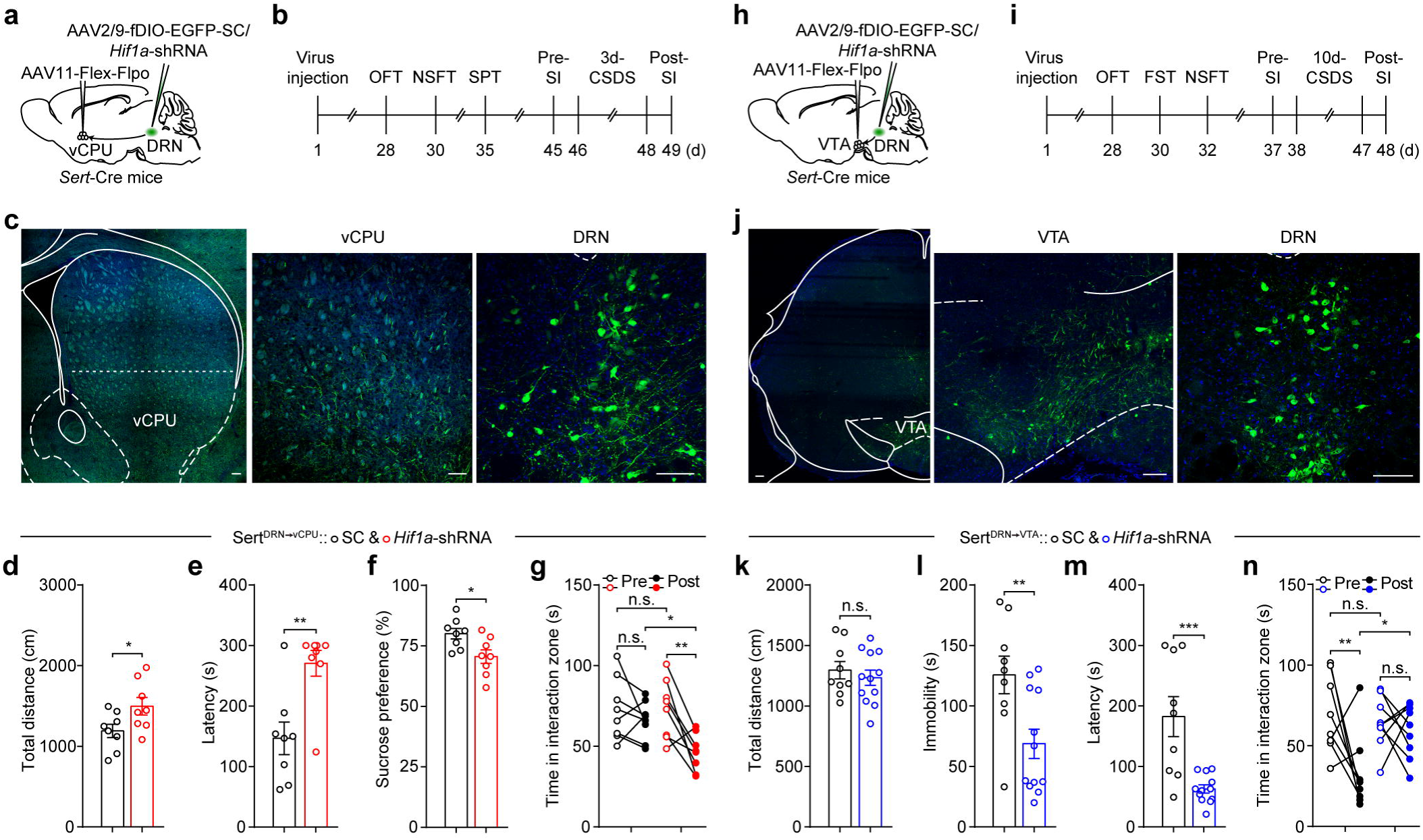
DRN^5-HT^ neurons differentially regulate depression-related behaviors via the HOSP. **a, h.** Flex-flpo strategy for knocking down *Hif1a* in Sert^DRN→vCPU^ (**a**) (*n* = 8) and Sert^DRN→VTA^ (**h**) (*n*_SC_ = 8–9; *n_Hif1a_*_-shRNA_ = 9–12) neurons. AAV11-Flex-Flpo virus was injected into the ventral caudal putamen (vCPU) or the ventral tegmental area (VTA) of *Sert*-Cre mice with infusion of AAV2/9-fDIO-*Hif1a*-shRNA or control virus with a scrambled sequence (SC) into the DRN. **b, i.** Experimental procedures for downregulating HIF1α in Sert^DRN→vCPU^ (**b**) and Sert^DRN→VTA^ (**i**) neurons. **c, j.** Confocal images of DRN inputs in the vCPU (**c**) and the VTA (**j**). **d–g, k–n.** Behavioral tests of knocking down *Hif1a* in Sert^DRN→vCPU^ (**d–g**) (*n* = 8) and Sert^DRN→VTA^ (**k–n**) (*n*_SC_ = 8–9; *n_Hif1a_*_-shRNA_ = 9–12) neurons. Total distance traveled in the OFT (**d, k**), latency in the NSFT (**e, m**), SPT ratio (**f**), immobility in the FST (**l**), and social interaction test after 3-(**g**) or 10-day (**n**) CSDS. Scale bars, 100 μm; \**p* < 0.05, \*\**p* < 0.01, and \*\*\**p* < 0.001. **d, e, f, k, l, m,** two-tailed unpaired *t*-test. **g, n,** two-way ANOVA with Sidak’s multiple comparisons test. n.s., not significant. Data are presented as mean ± SEM.

To further characterize DRN^5-HT^ neurons by their individual responses to stress via the HOSP, we expressed *Hif1a*-shRNA in Sert^DRN→vCPU^ (DRN^5-HT^ neurons projecting to the vCPU) or Sert^DRN→VTA^ (DRN^5-HT^ neurons projecting to the VTA) neurons in *Sert*-Cre mice. Behavioral analysis revealed that knocking down HIF1α in Sert^DRN→vCPU^ neurons induced a depressive phenotype, as evidenced by increased latency in the NSFT, a decreased sucrose preference ratio, and reduced social time in the interaction zone following a 3-day CSDS paradigm. Notably, expression of *Hif1a*-shRNA in Sert^DRN→VTA^ neurons had antidepressant-like effects, decreasing the duration of immobility in the FST, reducing latency in the NSFT, and abolishing social avoidance behaviors induced by a 10-day CSDS paradigm. We found that knocking down *Hif1a* in Sert^DRN→vCPU^ neurons increased locomotor activity in the OFT, whereas knocking down *Hif1a* in Sert^DRN→VTA^ neurons did not (Fig. 5a–5n). In addition, compared with controls, expression of *Hif1a*-shRNA in Sert^DRN→VTA^ or Sert^DRN→vCPU^ neurons had little effect on anxiety-like behaviors or cognition (Supplementary Fig. 6a–6j). Taken together, these results clearly show that the HOSP has opposite effects on Sert^DRN→VTA^ and Sert^DRN→vCPU^ neurons, leading to either the prevention or development of depressive behaviors.

**Fig. 6:**
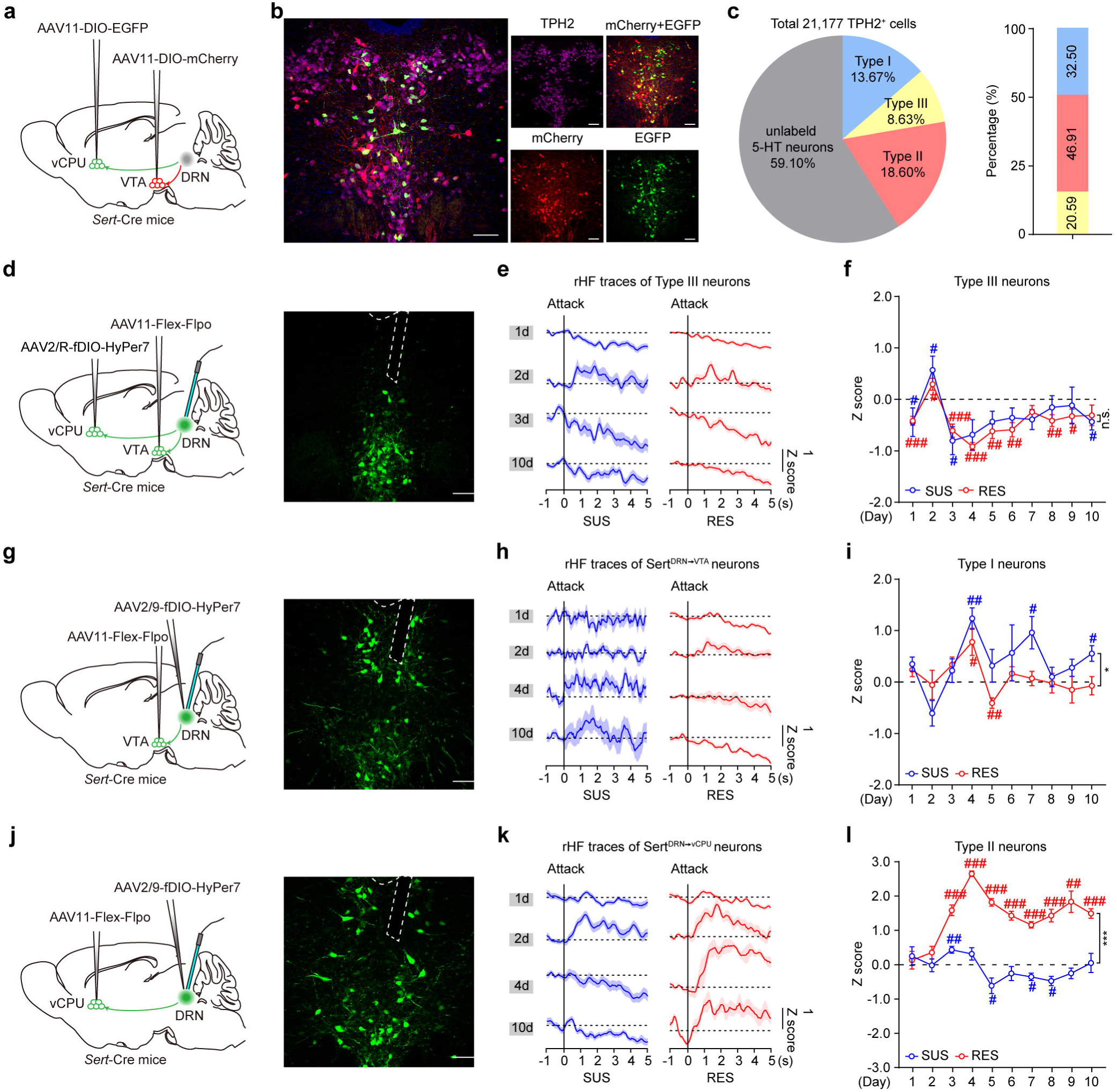
DRN^5-HT^ neurons scale O₂ utilization in response to traumatic stress. **a.** Strategy for labeling DRN^5-HT^ neurons projecting to the VTA and the vCPU. AAV11-DIO-EGFP was bilaterally injected into the vCPU of *Sert*-Cre mice, and AAV11-DIO-mCherry was injected into both sides of the VTA. **b.** Images of retrogradely labeled DRN^5-HT^ neurons. Violet, anti-TPH2 antibody; mCherry⁺ cells, Sert^DRN→VTA^ neurons; EGFP⁺ cells, Sert^DRN→vCPU^ neurons. **c.** Percentages of three types of DRN^5-HT^ neurons in the DRN (*n* = 7 mice). **d.** HyPer7 recordings of type III neurons. Strategy for labeling type III neurons and representative confocal image showing that type III neurons mainly locate in the ventral part of the DRN. **e.** Attack-locked rHF traces of type III neurons in response to a 10-day CSDS. The black lines indicate the onset of attacks. **f.** Mean attack-locked rHF of type III neurons in SUS and RES mice during a 10-day CSDS paradigm (*n* = 6/12 in SUS/RES mice, respectively). **g.** HyPer7 recordings of Sert^DRN→VTA^ neurons. **h.** Attack-locked rHF traces of Sert^DRN→VTA^ neurons following a 10-day CSDS. **i.** Mean attack-locked rHF of type I neurons in SUS and RES mice (*n* = 5/8 in SUS/RES mice). **j.** HyPer7 recordings of Sert^DRN→vCPU^ neurons. **k.** Attack-locked rHF traces of Sert^DRN→vCPU^ neurons following a 10-day CSDS. **l.** Mean attack-locked rHF of type II neurons in SUS and RES mice (*n* = 6/6 in SUS/RES mice). Scale bars, 100 μm; \**p* < 0.05 and \*\*\**p* < 0.001. **f, i, l,** two-way RM ANOVA with Sidak’s multiple comparisons test. ^#^*p* < 0.05, ^#^ ^#^*p* < 0.01, and ^#^ ^#^ ^#^*p* < 0.001 relative to the basal level (dashed line) by one-sample *t*-test (**f, i, l**). n.s., not significant. Data are presented as mean ± SEM.

Given that the dopaminergic mesocorticolimbic pathway is implicated in emotion regulation ^28,38,39^ and that DRN^5-HT^ neurons send excitatory inputs to dopamine neurons in the VTA that project to the nucleus accumbens (NAc) ^40^, we propose that inhibition of dopaminergic neurons in the mesoaccumbens projection could contribute to the antidepressant effect of *Hif1a* knockdown in Sert^DRN→VTA^ neurons. Using whole-cell electrophysiological recording, we found that knocking down *Hif1a* in Sert^DRN→VTA^ and Sert^DRN→vCPU^ neurons significantly decreased their firing rates, indicating that the NOX4–HOSP axis regulates the activities of these neurons and thereby modulates the related neural circuits. Notably, the firing rate of neurons in the mesoaccumbens projection was significantly lower in mice with *Hif1a* knockdown in Sert^DRN→VTA^ neurons than in control mice, indicating inhibition of these neurons (Supplementary Fig. 7a–7c). These results are consistent with previous reports ^28,38,39^, further supporting the notion that DRN^5-HT^ neurons differentially regulate depression-related behaviors via the HOSP.

**Fig. 7:**
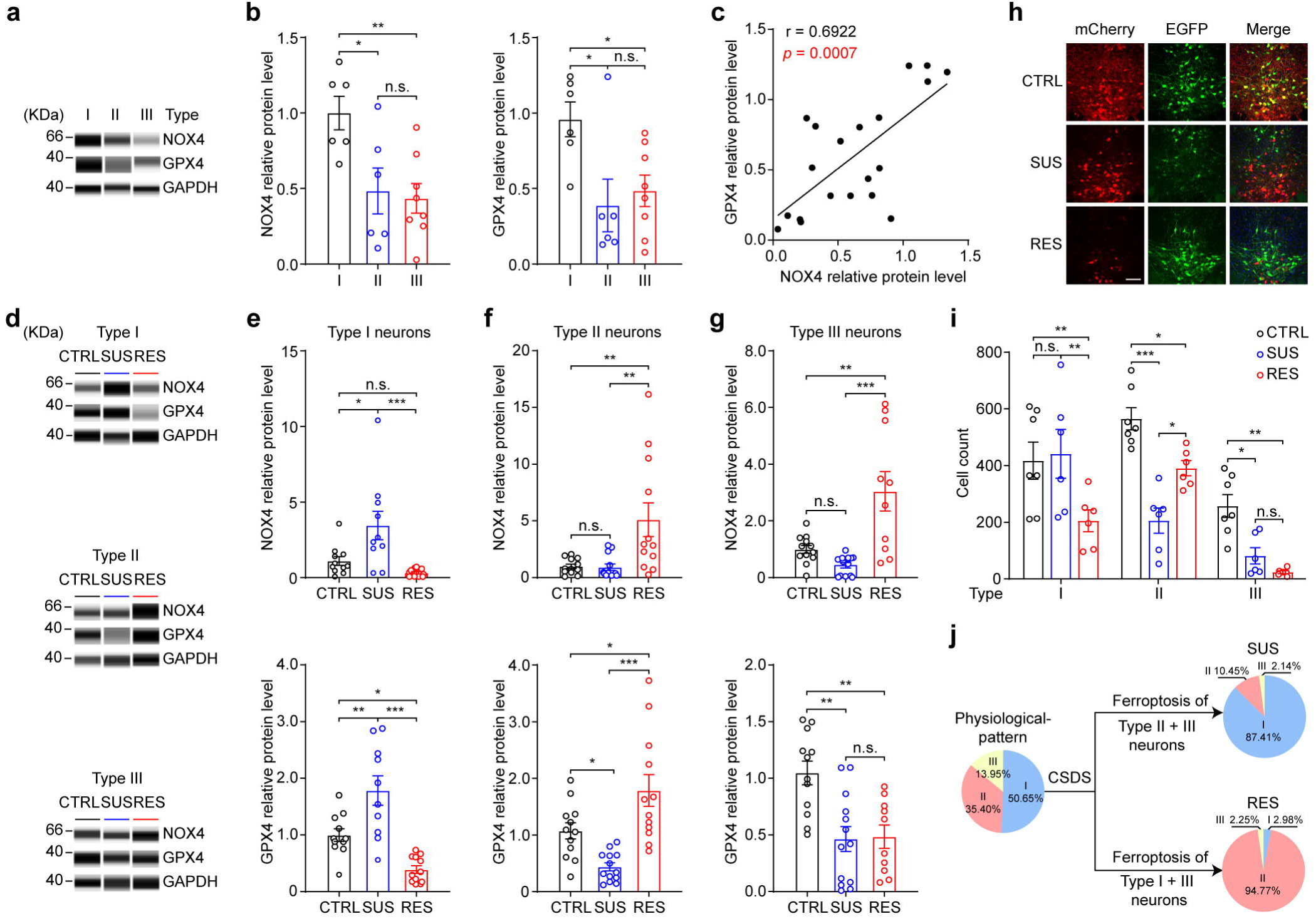
Ferroptosis mediates the scaling of O₂ utilization in DRN^5-HT^ neurons following CSDS. **a, b.** Simple western blot images and quantification of the levels of NOX4 and GPX4 in DRN^5-HT^ neurons projecting to the VTA and the vCPU under physiological conditions (*n*_type-I_ = 6, *n*_type-II_ = 6, *n*_type-III_ = 8 cells from 3–4 mice). **c.** Correlation between NOX4 expression and GPX4 protein levels under physiological conditions. Pearson’s correlation test. **d.** Representative images of simple western blots showing NOX4 and GPX4 expression in three types of DRN^5-HT^ neurons following a 10-day CSDS paradigm. **e–g.** Quantification of NOX4 and GPX4 for type I (**e**), type II (**f**), and type III (**g**) neurons (*n* = 10–13 cells from 3–4 mice per phenotype), respectively. h. Representative images of retrogradely labeled DRN^5-HT^ neurons following a 10-day CSDS paradigm. mCherry⁺ cells, type I neurons; EGFP⁺ cells, type II neurons; yellow (mCherry⁺ and EGFP⁺) cells, type III neurons. i. Quantification of three types of labeled DRN^5-HT^ neurons (*n*_CTRL_ = 7, *n*_SUS_ = 6, *n*_RES_ = 6 mice). j. Schematic diagram showing two patterns of O₂ scaling among DRN^5-HT^ neurons following CSDS. Scale bars, 100 μm; \**p* < 0.05, \*\**p* < 0.01, and \*\*\**p* < 0.001. **b, e, f, g,** one-way ANOVA with Tukey’s multiple comparisons test. **i,** two-way ANOVA with Bonferroni’s multiple comparisons test. n.s., not significant. Data are presented as mean ± SEM.

### DRN^5-HT^ neurons scale O_2_ utilization in response to traumatic stress

To define how changes in O₂ utilization in DRN^5-HT^ neurons affect behavioral outcomes in response to trauma, we characterized Sert^DRN→VTA^ and Sert^DRN→vCPU^ neurons. To this end, we injected the retrovirus AAV11-DIO-EGFP into the vCPU and the AAV11-DIO-mCherry retrovirus into the VTA of *Sert*-Cre mice to label these DRN^5-HT^ neurons (Fig. 6a and Supplementary Fig. 8a). We analyzed a total of 21,177 TPH2⁺ (a marker for DRN^5-HT^ neurons ^31^) cells in the DRN from seven brains and found that approximately 41% of TPH2⁺ neurons were labeled, divided into three types: type I neurons, mCherry⁺ only, indicating projection exclusively to the VTA; type II neurons, EGFP⁺ only, indicating projection exclusively to the vCPU; and type III neurons, mCherry⁺ and EGFP⁺, indicating projection to both the VTA and the vCPU (Fig. 6b and 6c).

To characterize dynamic changes in O₂ consumption by these DRN^5-HT^ neurons in mice subjected to a 10-day CSDS paradigm, we monitored intracellular H₂O₂ content in real time using the genetically encoded fluorescent H₂O₂ sensor HyPer7, a monomeric, ratiometric probe that specifically and reversibly senses H₂O₂ with high temporal resolution ^41^. We found that the ratio of <u>H</u>yPer7 fluorescence intensity (rHF) responded dose-dependently to H₂O₂ treatment in cultured HEK293 cells transfected with AAV2/9-CMV-Cre and AAV2/9-DIO-HyPer7, whereas EGFP-expressing HEK293 cells did not respond to H₂O₂ application, validating the HyPer7 recording method. We then injected AAV2/9-CMV-Cre and AAV2/9-DIO-HyPer7 into the DRN of adult C57BL/6J mice and recorded HyPer7 fluorescence intensity in the DRN during three distinct social behaviors in CSDS: attack, approach, and non-approach ^42^. We found that changes in rHF corresponded with attack bouts but not with approach or non-approach behavior (Supplementary Fig. 8b–8f), indicating that changes in O₂ utilization in the DRN are attack-locked events.

**Fig. 8:**
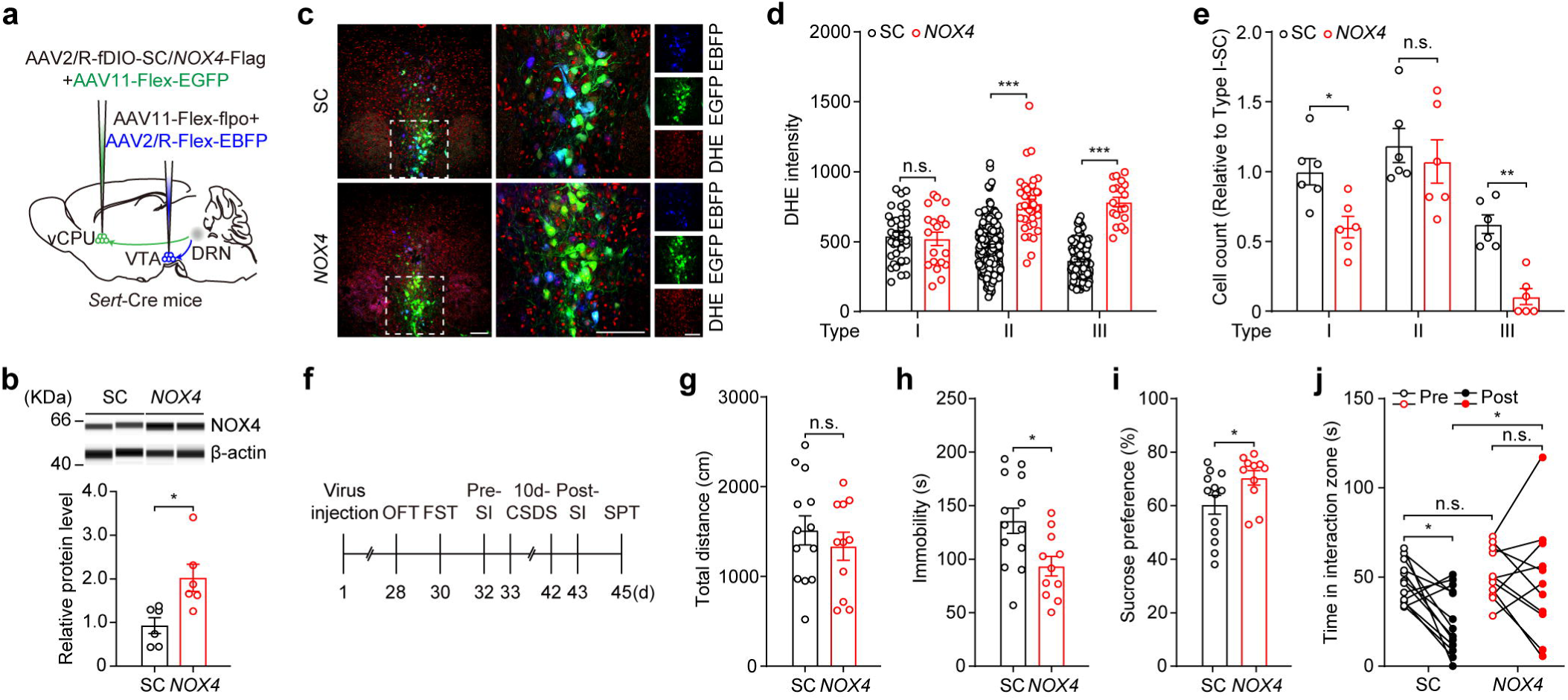
Increased O₂ consumption in type III neurons induces a resilient phenotype. **a.** Schematic of *NOX4* overexpression in type III neurons of *Sert*-Cre mice. **b.** Protein levels of NOX4 in the DRN of mice with *NOX4* expression in type III neurons and control mice (SC, scrambled sequence) 28 days after virus injection (*n* = 6). **c.** Representative images of virus-infected DRN^5-HT^ neurons and DHE fluorescence (red); green, Sert^DRN→vCPU^ neurons; blue, Sert^DRN→VTA^ neurons. **d.** Quantification of cellular DHE fluorescence intensity in the three types of DRN^5-HT^ neurons (*n* = 19–40 cells for type I neurons, *n* = 39–187 cells for type II neurons, *n* = 19–137 cells for type III neurons, respectively, from four mice per group). **e.** Numbers of the three types of DRN^5-HT^ neurons in NOX4-overexpression mice (*NOX4*) and control mice (SC) (*n* = 6). **f.** Behavioral tests for mice with NOX4 overexpression in type III neurons and control mice injected with AAV2/R-fDIO-SC into the vCPU. **g–j.** Effects of NOX4 overexpression in type III neurons on locomotion in the OFT (**g**), immobility in the FST (**h**), SPT ratio (**i**), and time spent in the interaction zone following a 10-day CSDS paradigm (**j**) compared with those of control animals (*n*_SC_ = 13, *n_NOX4_* = 11). Scale bars, 100 μm; \**p* < 0.05, \*\**p* < 0.01, and \*\*\**p* < 0.001. **b, g, h, I,** two-tailed unpaired *t*-test. **d, e,** two-way ANOVA with Bonferroni’s multiple comparisons test. **j,** two-way ANOVA with Sidak’s multiple comparisons test. n.s., not significant. Data are presented as mean ± SEM.

We subsequently investigated dynamic changes in O₂ consumption by DRN^5-HT^ neurons using fluorescence imaging with the HyPer7 probe. To label type III neurons, we applied a Flex-Flpo strategy in which retrograde AAV2/R-fDIO-HyPer7 virus was microinjected into the vCPU and AAV11-Flex-Flpo virus into the VTA of *Sert*-Cre mice (Fig. 6d). Four weeks after virus injection, mice were subjected to a 10-day CSDS paradigm, and HyPer7 fluorescence intensity was recorded. We found that the rHF of type III neurons (hereafter referred to as rHF^type-III^) during attack bouts dramatically fluctuated during the early stage of CSDS—decreasing on the first day, reaching its highest level on the second day, and then decreasing to its lowest level on the third day of the CSDS paradigm (Fig. 6e and 6f). However, there was no significant difference between RES and SUS mice.

To characterize changes in rHF in type I neurons (rHF^type-I^), we recorded the HyPer7 fluorescence intensity of Sert^DRN→VTA^ neurons (Fig. 6g), and rHF^type-I^ was calculated as the rHF of Sert^DRN→VTA^ neurons minus that of rHF^type-III^ (see details in the Methods). From the first to the third day of CSDS, attack-locked rHF^type-I^ in SUS mice fluctuated, whereas in RES mice, it remained stable relative to the basal level. Moreover, rHF^type-I^ fluctuated during attack bouts from the fourth to the tenth day of CSDS, but displayed opposite trends in SUS and RES mice (Fig. 6h, 6i and Supplementary Fig. 8g). Similarly, changes in rHF of type II neurons (rHF^type-II^ = rHF of Sert^DRN→vCPU^ − rHF^type-III^) corresponded to attack periods during the 10-day CSDS paradigm. rHF^type-II^ did not change on the first day of CSDS in either RES or SUS mice but increased on the third day, showing a significant difference between the two groups. Notably, after the third day of CSDS, rHF^type-II^ changed in opposite directions in RES and SUS mice during attacks. In RES mice, rHF^type-II^ further increased and remained stable at a high level, whereas in SUS mice, it decreased to the basal level and remained stable below baseline (Fig. 6j–6l, Supplementary Fig. 8h and 9a–9c).

These behavior-related changes in rHF strongly suggest that cellular O₂ utilization in DRN^5-HT^ neurons projecting to the VTA and vCPU follows two distinct patterns, referred to here as the RES and SUS patterns, respectively, in response to attacks during the CSDS paradigm. Moreover, we observed that RES and SUS patterns, reflected by changes in rHF in DRN^5-HT^ neurons, correlated with behavioral outcomes following the SEFL paradigm (Supplementary Fig. 9d–9f). In addition, we observed no changes in rHF in these DRN^5-HT^ neurons in mice subjected to a chronic restraint stress paradigm or to a series of mechanical pain stimuli (Supplementary Fig. 9g–9j). Notably, only rHF^type-III^ was altered in response to the first attack in the CSDS paradigm, indicating that type III neurons are the first to respond to traumatic stress (Supplementary Fig. 9k–9l). Taken together, these data indicate that DRN^5-HT^ neurons scale O₂ utilization in response to traumatic stress, thereby affecting behavioral outcomes.

### Ferroptosis mediates the scaling of O_2_ utilization in DRN^5-HT^ neurons following CSDS

To investigate the molecular mechanism by which DRN^5-HT^ neurons scale O₂ utilization in response to trauma, we developed a method for analyzing single-cell protein expression under low *p*O₂ conditions that mimic physiological environments. We found that protein levels of NOX4 in type I neurons were the highest among DRN^5-HT^ neurons, while NOX4 expression in type II neurons was equal to that in type III neurons (Fig. 7a and 7b), indicating that O₂ utilization in DRN^5-HT^ neurons projecting to the VTA and vCPU is heterogeneous under physiological conditions.

To characterize the effect of trauma on O₂ utilization in DRN^5-HT^ neurons, we examined NOX4 expression in these neurons following a 10-day CSDS paradigm. We found that NOX4 levels in type I neurons were significantly increased in SUS mice compared with controls, but not in RES mice. However, in both type II and III neurons, NOX4 expression dramatically increased—by approximately fivefold and threefold, respectively—in RES mice compared with controls. No changes in NOX4 protein levels were observed in type II and III neurons from SUS mice (Fig. 7d–7g). These data clearly show that distinct subpopulations of DRN^5-HT^ neurons projecting to the VTA and vCPU differentially regulate O₂ utilization in response to traumatic stress.

Clinical studies have shown that the number of neurons decreases in the ventral part of the DRN (vDRN) ^43^, and both NOX4 and H₂O₂ contribute to ferroptosis ^44^. Here, we observed that type III neurons were located mainly in the vDRN (Fig. 6d) and that both rHF and NOX4 expression of DRN^5-HT^ neuron subtypes changed dramatically—especially rHF^type-III^—during the early stage of CSDS when behavioral outcomes were determined. Therefore, we proposed that ferroptosis contributes to the altered pattern of O₂ utilization in DRN^5-HT^ neurons following CSDS. Supporting this hypothesis, we found that the number of TUNEL⁺ cells was dramatically greater in the DRN following a 10-day CSDS paradigm in model animals than in controls. However, expression of caspase-3, a marker of apoptosis ^45^, was unaltered in the DRN. Notably, the number of TPH2⁺ cells decreased to a similar extent in both SUS and RES mice compared with controls (Supplementary Fig. 10a–10e), indicating that DRN cell death induced by traumatic stress was cell-type dependent. We then performed cell counting in the DRN following cell labeling and found that the number of type I neurons dramatically decreased in RES mice subjected to a 10-day CSDS paradigm compared with controls, with no changes observed in SUS mice. Although some type II neurons were lost in both RES and SUS mice, the number of surviving type II neurons was greater in RES mice than in SUS mice. Notably, the number of type III neurons significantly decreased following CSDS, but there was no difference between the two groups (Fig. 7h and 7i).

To further characterize ferroptosis of DRN^5-HT^ neurons following CSDS, we examined expression of glutathione peroxidase 4 (GPX4), the central regulator of ferroptosis ^44^. Under physiological conditions, similar to NOX4, GPX4 levels in type I neurons were the highest, and expression levels in type II and III neurons were equal (Fig. 7a and 7b). Moreover, there was a positive correlation between NOX4 and GPX4 expression among DRN^5-HT^ neurons (Fig. 7c), indicating that coordination of NOX4 and GPX4 is maintained under physiological conditions. Following the 10-day CSDS paradigm, GPX4 protein levels in type I neurons were increased in SUS mice but significantly decreased in RES mice compared with controls (Fig. 7d and 7e). In contrast, in type II neurons, GPX4 expression was downregulated in SUS mice but upregulated in RES mice compared with controls (Fig. 7d and 7f). Moreover, there was a positive correlation between GPX4 and NOX4 protein levels in type I neurons of SUS mice and in type II neurons of RES mice (Supplementary Fig. 10f and 10g), further supporting the notion that coordination of NOX4 and GPX4 determines the cell fate of DRN^5-HT^ neurons following CSDS. Notably, GPX4 expression was downregulated in type III neurons after the 10-day CSDS paradigm; however, no difference was found between the two mouse groups (Fig. 7d and 7g). These results clearly reveal that ferroptosis contributes to altered patterns of O₂ utilization in DRN^5-HT^ neurons following traumatic stress.

Based on NOX4 expression and neuronal survival, we calculated O₂ consumption in the three DRN^5-HT^ neuron subgroups. In RES mice, O₂ utilization increased in type II neurons and decreased in type I and III neurons compared with controls. However, in SUS mice, type I neurons consumed more O₂, whereas type II and III neurons consumed less O₂ than in controls (Fig. 7j). Notably, these behavior-related O₂ utilization patterns were consistent with those observed using HyPer7 fluorescence imaging on the tenth day of CSDS (Fig. 6f, 6i, and 6l), suggesting that altered O₂ utilization patterns in DRN^5-HT^ neurons projecting to the VTA and vCPU control behavioral outcomes in response to traumatic stress.

### Increased O₂ consumption in type III neurons induces a resilient phenotype

Because rHF^type-III^ fluctuated dramatically from the first to the third day of the CSDS paradigm, and the behavioral phenotype was first observed on the third day ^42^, we proposed that increased O₂ consumption in type III neurons triggers O₂ utilization scaling in the DRN. To test this hypothesis, we overexpressed *NOX4* in type III neurons, and 4 weeks after virus injection, prepared brain slices for DHE staining and neuronal quantification (Fig. 8a). We found that *NOX4* overexpression in type III neurons significantly increased NOX4 protein levels in the DRN of model mice compared with controls (Fig. 8b). DHE staining revealed that, compared with controls, DHE fluorescence intensities in both type II and III neurons were greater, while no significant difference was observed in type I neurons between the two groups (Fig. 8c and 8d). Moreover, the numbers of type I and III neurons were dramatically lower, whereas the number of type II neurons was unchanged in experimental mice compared with controls (Fig. 8e). Based on neuronal survival and cellular DHE intensity, we estimated that *NOX4* overexpression in type III neurons induced a RES-like pattern of O₂ utilization in DRN^5-HT^ neurons, with increased O₂ consumption in type II neurons and decreased O₂ consumption in type I and III neurons (Supplementary Fig. 10h).

We then used a new group of *Sert*-Cre mice, and behavioral tests were conducted 4 weeks after virus injection (Fig. 8f). Notably, behavioral testing revealed that, compared with controls, mice overexpressing *NOX4* in type III neurons exhibited a resilient phenotype, as evidenced by decreased immobility in the FST without affecting locomotion in the OFT, an increased sucrose preference ratio, and increased time spent in the interaction zone when CD1 mice were present following a 10-day CSDS paradigm (Fig. 8g–8j). Together, these results strongly suggest that ferroptosis in DRN^5-HT^ neurons projecting to the VTA and vCPU regulates patterns of O₂ utilization, which in turn control behavioral outcomes in response to traumatic stress.

## DISCUSSION

An adequate supply of O₂ is essential for mitochondrial respiration and oxidative phosphorylation, making it crucial for the survival of all metazoans ^46,47^. However, within the brain, the *p*O₂ is as low as 1–40 mmHg, and the role of such low and heterogeneous *p*O₂ remains unknown. Here, we identified a novel function of O₂ in the brain, in which O₂ actively initiates cell signaling cascades that control behavioral outcomes in response to traumatic stress.

Some trauma-exposed individuals maintain mental health, whereas others develop depressive disorders. How does trauma induce different behavioral outcomes over a short timescale? We showed that O₂ in the DRN plays a key role in this process. DRN^5-HT^ neurons projecting to the VTA and vCPU differentially respond to traumatic stress, such as attacks in the CSDS paradigm, by scaling O₂ utilization in two distinct patterns—the RES and SUS patterns—and thereby modulating corresponding behaviors. In contrast, chronic restraint stress and mechanical pain stimulation cannot induce such O₂ utilization scaling in the DRN, possibly because these stresses are not as etiologically relevant as social stress^4^.

How does O₂ utilization in DRN^5-HT^ neurons regulate behavior? The answer is that DRN^5-HT^ neurons projecting to the VTA and vCPU differentially regulate depression-related behaviors via the NOX4–HOSP axis (Supplementary Fig. 11). Approximately 41% of DRN^5-HT^ neurons project to the VTA and vCPU brain regions, and these neurons can be divided into three types—type I, II, and III—whose activity is regulated by the NOX4–HOSP axis. Under physiological conditions, type I neurons express the highest level of NOX4 and consume more O₂ than type II and III neurons. Because GPX4 expression is increased in type I neurons, these neurons survive.

In response to traumatic stress, type III neurons (8.63% of DRN^5-HT^ neurons) respond first, as evidenced by increased NOX4 expression and upregulation of rHF. However, GPX4 expression in type III neurons cannot be correspondingly upregulated, and thus most of this subgroup dies. This explains why rHF^type-III^ fluctuated dramatically during the early stage of CSDS and remained at a lower level relative to baseline during the later stage. Moreover, NOX4 overexpression in type III neurons induced ferroptosis of type I and III neurons, indicating that increased O₂ utilization in type III neurons triggered O₂ redistribution within the DRN. Notably, we found that NOX4 overexpression in type III neurons induced only a RES-like pattern of O₂ utilization and resilient behaviors, but not depression. This may be because certain DRN^5-HT^ neurons (e.g., type III and II) intrinsically sense O₂ increases, as they exhibit low basal O₂ utilization and GPX4 levels. In contrast, type I neurons utilize more O₂ and express higher GPX4 levels, making them more sensitive to O₂ deficiency rather than moderate increases in O₂ supply. Further experiments are needed to test this hypothesis.

After ferroptosis of type III neurons, O₂ in the DRN likely needs to be redistributed. In type I neurons, NOX4 expression was upregulated and O₂ consumption increased, but GPX4 expression did not correspondingly increase; thus, some of these neurons died. Meanwhile, type II neurons consumed more O₂ and coordinately increased GPX4 expression, resulting in a RES pattern of O₂ utilization. Consistently, rHF^type-II^ further increased during the third and fourth days of CSDS and remained at a high level thereafter, whereas rHF^type-I^ dramatically decreased during this period and remained low during the later stage of CSDS. Type I neurons project excitatory inputs onto VTA dopaminergic neurons that project to the NAc ^40^; therefore, under the RES pattern of O₂ utilization, the death of some type I neurons reduced the activity of dopaminergic neurons in the mesoaccumbens projection, which has been implicated in the development of resilient behavior following CSDS ^39^.

When type I neurons upregulated GPX4 expression in proportion to increased O₂ consumption, they survived, and their neuronal activity increased. Moreover, GPX4 expression was downregulated in type II neurons, and some of these neurons died, suggesting a SUS pattern of O₂ utilization, in which rHF^type-II^ decreased dramatically from the third to fourth days of the CSDS paradigm and remained low thereafter, whereas rHF^type-I^ fluctuated but remained above baseline. Under the SUS pattern of O₂ utilization, increased activity of type I neurons activated mesoaccumbens-projecting dopaminergic neurons in the VTA, which has been implicated in the development of the SUS phenotype in mice following a 10-day CSDS paradigm ^28,39^.

In summary, we found that O₂ utilization by DRN^5-HT^ neurons projecting to the VTA and vCPU is heterogeneous under physiological conditions. In response to traumatic stress, these neurons scale O₂ utilization in two distinct patterns that control behavioral outcomes. These results highlight that an in-depth understanding of O₂ regulation mechanisms in the brain could open new avenues for the treatment and prevention of stress-related disorders, particularly PTSD.

## METHODS

### Mice

All animal procedures were performed in accordance with the institutional guidelines of South China University of Technology (Guangzhou, China) and complied with the Guidelines for Animal Experimentation of the Chinese Council Institutes. The following mouse strains were used: *Sert*-Cre (MMRRC stock 031028-UCD), *Gad2*-Cre (JAX stock 028867). All transgenic mice were back-crossed to C57BL/6J background. Male C57BL/6J mice (aged 8–10 weeks) and male CD1 mice (aged 8–9 months) were purchased from the Beijing Vital River Laboratory Animal Technology Co., Ltd (Beijing, China). All mice were maintained under a 12-h light–dark cycle with *ad libitum* access to food and water.

### Human brain samples

This study was approved by the Douglas Hospital Research Ethics Board. Postmortem brain samples were provided by the Suicide Section of the Douglas-Bell Canada Brain Bank (http://www.douglas.qc.ca/page/brain-bank). These included samples from subjects who died by suicide in the context of a depressive episode and samples from matched controls who died accidentally or of natural causes and had no history of psychiatric, neurological, or inflammatory illnesses. Unfixed frozen tissue samples containing the dorsal raphe nucleus (DRN) were dissected and stored at –80°C until used. The groups were matched by age, tissue pH, and postmortem interval. Detailed subject information is provided in **Supplementary Table 1**.

### Dihydroethidium (DHE) staining

Mice were anesthetized with pentobarbital sodium (75mg/kg, intraperitoneal injection) and whole brains were collected and placed in room-temperature PBS prior to embedding in Tissue-Tek OCT Compound (Sakura Finetek, Torrance, CA, USA). Embedded tissues were immediately placed on dry ice and sliced into 40-μm-thick sections using a freezing microtome (CM1950; Leica Microsystems, Wetzlar, Germany). For sagittal sections, slices spanning the entire mediolateral axis were systematically collected. Every third slice was retained, with a total of three sets of brain slices being collected. Preparations of coronal sections containing the DRN followed precise neuroanatomical protocols. Perpendicular coronal cuts were made to isolate the midbrain region containing the DRN (AP −4.12 to −4.84 mm from bregma) according to The Mouse Brain in Stereotaxic Coordinates (2001, second edition) by George Paxinos & Keith B. J, Franklin ^48^. Serial sections (40 μm) were obtained using a cryostat, with every fourth section being retained, yielding four sets of brain slices.

For DHE staining, sections (40 μm) were mounted on microscope glass slides and incubated for 30 min at 37°C with 10 μM DHE (D7008; Sigma-Aldrich, St. Louis, MO, USA). After washing three times with PBS, 5 min each wash, the sections were mounted with mounting medium (Vectashield; Vector Laboratories, Burlingame, CA, USA), and imaged using an A1R confocal microscope (Nikon Instruments, Tokyo, Japan).

Mean DHE fluorescence intensity in each brain area was quantified using Imaris x64 9.7.1 software (Oxford Instruments, Oxford, England). High-resolution confocal z-stacks were imported into Imaris and anatomical regions were semi-automatically segmented using the Surface Creation module. Region-specific fluorescence intensity was quantified by applying intensity thresholding and voxel-based measurements, with background subtraction performed using adjacent non-fluorescent areas as reference. The intensities and areas of the brain regions were input into the software’s integrated Statistics module and the mean DHE fluorescence intensity was calculated by dividing the total fluorescence intensity by the total area of the corresponding region.

NIS-Elements analysis software (5.21, Nikon) was used for the quantification of DHE fluorescence intensity in individual cells. Cells were segmented using the ROI module *via* adaptive thresholding based on the expression of fluorescence markers such as EGFP and EBFP. Fluorescence intensity (TRITC channel) was calculated as integrated density normalized to cell volume.

### Optical fiber/cannula implantation

Mice were anesthetized and then placed on a stereotaxic frame (RWD Life Science, Shenzhen, China). Adeno-associated viruses (AAVs) (viruses are listed in **Supplementary Table 2**) were injected through a micro-syringe pump (Stoelting, Wood Dale, IL, USA) fitted with a 33-gauge needle at a rate of 0.1 µL min^−1^. After injection, the cannula was left *in situ* for 10 min and then slowly removed.

The injection coordinates and volumes (mediolateral [ML], anteroposterior [AP], and dorsoventral [DV] coordinates relative to the bregma) used were:

mPFC: ML = ± 0.35 mm, AP = +1.75 mm, DV = –2.50 mm; inject volume: 200 nL/site; NAc: ML = ± 0.75 mm, AP = + 1.50 mm, DV = – 4.50 mm; inject volume: 200 nL/site; dCPU: ML = ± 1.60 mm, AP = + 0.50 mm, DV = – 2.70 mm; inject volume: 200 nL/site; vCPU: ML = ± 2.00 mm, AP = + 0.50 mm, DV = – 3.50 mm; inject volume: 200 nL/site; VTA: ML = ± 0.35 mm, AP = - 3.30 mm, DV = – 4.25 mm; inject volume: 200 nL/site; MGN: ML = ± 2.00 mm, AP = - 3.20 mm, DV = – 2.80 mm; inject volume: 200 nL/site; DRN: ML = + 0.83 mm, AP = - 4.60 mm, DV = – 3.18 mm; inject volume: 500 nL (a 15° angle from caudal to rostral)

For the fiber photometry experiments, after virus injection, a ceramic ferrule (200 µm; Inper, Hangzhou, China) was surgically implanted above the DRN of each mouse. Mice were allowed to recover for at least 5 days before recordings.

For the infusion of FG-4592 (S1007; Selleck Chemicals, Houston, TX, USA), a unilateral cannula (RWD Life Science) was implanted above the DRN and secured with dental cement. A stainless-steel obturator was inserted into each guide cannula to prevent blockage. Five days following cannula implantation, mice received 1 mL infusions of FG-4592 (2.5, 20, 50 μM) or vehicle at a rate of 0.1 mL/min with a syringe pump (RWD), and the infusion cannula was left *in situ* for additional 10 min. Then, mice were returned to their home cage, and behavioral tests were conducted 1 h later. Only mice with correctly positioned optical fibers/cannula and exhibiting viral expression in the correct locations were used for further analysis.

### Fiber photometry

Following the injection of AAV2/9-DIO-HyPer7, AAV2/9-fDIO-HyPer7, or AAV2/R-fDIO-HyPer7, a ceramic optical fiber ferrule (200 μm diameter, 0.37 numerical aperture [NA]; Inper) was implanted with the fiber tip positioned in the DRN of *Sert*-Cre and *Gad2*-Cre mice. Photometric recordings were conducted using a fiber photometry recording system (Inper) 3 weeks after optical fiber implantation. To record fluorescence signals, two laser beams (410, reference light source and 499 nm, light source excited HyPer7) were reflected off a dichroic mirror, focused by a 10 × objective (NA 0.3), and then channeled through an optical commutator to the sample. The resulting fluorescence was bandpass filtered (MF525-39; Thorlabs, Newton, NJ, USA) and collected by a photomultiplier tube (R3896; Hamamatsu Photonics, Hamamatsu City, Japan). An amplifier (C7319; Hamamatsu Photonics) was used to convert the photomultiplier tube current output to voltage (V) signals, which were then further filtered through a low-pass filter (40-Hz cutoff; Brownlee 440; NeuroPhase, Santa Clara, CA, USA). To minimize photobleaching, the laser power at the fiber tip was adjusted to 30 μW. For recordings in mice during the chronic social defeat stress (CSDS), stress-enhanced fear learning (SEFL), restraint stress, and mechanical pain stimulating paradigms, fluorescence signals were acquired through optical fibers connected to the fiber photometry system. Bulk fluorescence signals were acquired and analyzed with MATLAB software. The F499/F410 ratio (rHF) was used to determine V_basal_ (baseline reference, 1 s pre-event) and V_Signal_ (signal during the event) and the z-score was calculated using the formula z-score = (V_signal_ –F_0_)/σ_F_, where F_0_ is the average value of V_basal_ and σ_F_ is the standard deviation of V_basal_. The mean z-score is the average value of z-score at each time point.

The mean rHF z-scores in type I and II neurons were calculated by a formula: the rHF z-score of the Sert^DRN→VTA^ and Sert^DRN→vCPU^ neurons minus the mean rHF z-score of type III neurons at each corresponding time point, respectively.

### Cell culture

To assess the sensitivity of HyPer7, fluorescence intensity was measured at 410 and 488 nm in response to varying H_2_O_2_ concentrations using the A1R confocal microscope (Nikon Instruments). Briefly, HEK293 cells were cultured in DMEM F12 (10-092-CV; Corning Inc, Corning, NY, USA) supplemented with 10% (*v*/*v*) FBS (16140089; Thermo Fisher) at 37°C in a 5% CO_2_ atmosphere. Subsequently, the cells were seeded in 35-mm glass-bottom dishes, and, after 24 h, they were transfected with CMV-Cre and AAV2/9-DIO-HyPer7 viruses or control virus AAV2/9-DIO-EGFP. For imaging experiments, 24 h after transfection, the culture medium was replaced with 1.2 mL of HBSS solution (H1387; Sigma-Aldrich) supplemented with 20 mM HEPES (H4034; Sigma-Aldrich). Cells were incubated in medium containing different concentrations of H_2_O_2_ (2, 100, 200, and 1000 μM), and real-time fluorescence was monitored using excitation wavelengths of 410 and 488 nm until a steady state had been reached. The HyPer7 ratio-metric signal was calculated by dividing the emission intensity obtained at 488 nm by that acquired at 410 nm.

### Cell Counting Kit–8 (CCK–8) assay

HEK293 cells were seeded in 96-well plates at a density of approximately 1 × 10^4^ cells/well. After culturing for 24 h at 37°C with 5% CO_2_, FG-4592 or vehicle was added to the cells. After an additional 48 h, 10 µL of CCK-8 reagent (C0037, Beyotime, Shanghai, China) was added to each well, and the plates were incubated for 1 h under the same conditions. Absorbance at 450 nm was measured using a microplate reader (PerkinElmer, Wellesley, MA, USA). Each group included at least three repetitions.

### NADH/NAD^+^ analysis

The effect of *Lb*NOX on modulating cytosolic O_2_ level was evaluated by measuring the NADH/NAD^+^ ratio as previously described, with minor modifications ^34^. Briefly, brain samples were extracted using a 4:4:2 mixture of acetonitrile/methanol/water containing 0.1 M formic acid and 5 µL of the resulting supernatant was injected onto a ZIC-pHILIC column (EE20000778; Millipore-Sigma). The column temperature was maintained at 27°C. Mobile phase A was 20 mM ammonium carbonate in water, pH 9.6 (adjusted with ammonium hydroxide), and mobile phase B was acetonitrile. The chromatographic separation gradient was as follows (flow rate: 0.15 ml/min): 0 min, 80% B; 0.5 min, 80% B; 20.5 min, 20% B; 21.3 min, 20% B; 21.5 min, 80% B, with a 7.5-min column equilibration period. The monitoring ions were m/z 664.0 → 136.0 for NAD^+^, 666.0 → 514.0 for NADH. Data were manipulated with SCIEX OS (ExionLC, Triple Quad 6500, SCIEX, USA) and Analyst 1.7 software (SCIEX). Individual NADH (N8129, Sigma), NAD^+^ standards (N7004, Sigma) were dissolved in 50% methanol at a concentration of 200 ng/mL.

### Quantitative real-time PCR (qPCR)

Total RNA was extracted using RNAiso Reagent according to the manufacturer’s instructions (9109, TaKaRa Bio, Osaka, Japan) and was reverse-transcribed to cDNA using PrimeScript RT Master Mix (RR047A, TaKaRa Bio). qPCR was performed with TB Green qPCR Master Mix (RR420A, TaKaRa Bio). The 18S rRNA gene was used as an internal control for normalization. Details of all the primers used in this study are provided in **Supplementary Table 3.**

### Western blot analysis

Tissue total protein was extracted using RIPA lysis buffer (P0013B; Beyotime) containing protease inhibitor (ST506-2; Beyotime). Cytoplasmic and nuclear fractions from brain tissues were separated using NE-PER Nuclear and Cytoplasmic Extraction Reagents (78833; Thermo Fisher Scientific). Protein concentrations in the resulting supernatant lysates were measured using a BCA kit (23225, Thermo Fisher Scientific). For human brain tissue, a traditional western blot protocol was employed, using anti-NOX4 (1:5000, 14347-1-AP; Proteintech, Rosemont, IL, USA) and anti-β-actin (1:2000, sc-1616; Santa Cruz, Santa Cruz, California, USA) primary antibodies. Protein levels were normalized to β-actin (for the cytoplasmic fractions) or Lamin B (for the nuclear fractions) on the same gel. Quantifications were performed with AlphaEase FC software (Alpha Innotech Corporation, USA).

For the Simple Western assay, the JESS system (Bio-Techne; ProteinSimple, San Jose, CA, USA) was used, employing the following primary antibodies: anti-HIF-1α (1:50, 14179S; Cell Signaling Technology, Danvers, MA, USA), anti-FIH1 (1:200, NB100-428SS; Novus Biologicals, Littelton, CO, USA), anti-PHD2 (1:200, NB100-2219, Novus Biological), anti-VHL (1:200, ab77262; Abcam, Cambridge, UK), anti-Lamin B (1:200, ab16048, Abcam), anti-NOX4 (1:500, 14347-1-AP; Proteintech), anti-caspase3 (1:500, 9662S; Cell Signaling Technology), anti-GPX4 (1:5000, ab125066; Abcam), and anti-β-actin (1:500, 20536-1-AP; Proteintech). Relative protein levels were determined *via* the peak areas detected in the chemiluminescence electropherogram generated by the Compass for SW software (ProteinSimple).

### HPLC-electrochemical detection (ECD) analysis

HPLC analysis was performed to determine the level of 5-HT and that of its metabolite 5-hydroxyindoleacetic acid (5-HIAA) in adult male *Sert*-Cre mice injected with AAV2/9-DIO-EGFP-*Hif1a*-shRNA or control virus into the DRN. Four weeks after virus injection, brain regions were collected and homogenized with 0.5 M perchloric acid, centrifuged, and the resulting supernatant was added to a microdialysis tube. To decrease the rate of 5-HT oxidation, the sample collection tubes were pretreated with an antioxidative agent (8 µL) containing 100 mM acetic acid (45754; Sigma-Aldrich), 0.2 mM disodium EDTA (E9884; Sigma-Aldrich), and 12.5 μM ascorbic acid (pH 3.2, A92902; Sigma-Aldrich).

The HPLC system consisted Nexera XR (SHIMADZU, Shimadzu, Japan), coupled with an amperometric electrochemical detector (DECADE Lite, Antec Scientific, Netherlands) in a wall-jet arrangement with a glassy carbon working electrode and an Ag/AgCl reference electrode in situ Ag/AgCl (ISAAC). Column C18 2.1× 100 mm, 3 µm (Sykam GmbH, Eresing, Germany) was used, and the mobile phase contained 8% methanol, sodium octane Sulfonate (0.160 g/L), sodium hydrogen phosphate (15.601 g/L), disodium ethylenediaminetetraacetate (8.835 mg/L), and potassium chloride (0.1492 g/L). The parameters for HPLC: total running time, 26.5 min; flow rate, 0.2 mL/min; the column oven temperature, 40℃; the detection potential, +0.54 V; the injection volume, 10 µL.

### Electrophysiological recording

Mice (8–14 weeks old) were deeply anesthetized and immediately decapitated. Then, the brains were rapidly extracted and dissected in ice-cold, oxygenated artificial cerebrospinal fluid (ACSF) composed of 195 mM sucrose, 2 mM KCl, 0.2 mM CaCl₂, 12 mM MgSO₄, 1.3 mM NaH₂PO₄, 26 mM NaHCO₃, and 10 mM D-glucose. Coronal brain sections (300-μm thick) containing the regions of interest were cut using a vibratome (VT-1200S, Leica, Germany). The slices were first incubated at 34°C for 30 min and subsequently allowed to recover at 25 ± 1°C in a holding chamber containing modified ACSF (126 mM NaCl, 26 mM NaHCO₃, 3 mM KCl, 1.25 mM NaH₂PO₄, 2 mM CaCl₂, 1 mM MgSO₄, 10 mM D-glucose). All solutions were continuously oxygenated with carbogen (95% O₂/5% CO₂, *v*/*v*). Electrophysiological recordings were conducted using EPC10 amplifiers (HEKA, Germany) in submerged brain slices perfused at a rate of 2 mL/min and maintained at 24 ± 2°C. The employed glass microelectrodes (4–6 MΩ resistance, pulled using a Narishige PC-100) were filled with a solution containing 130 mM K-D-gluconate, 20 mM KCl, 10 mM HEPES, 10 mM phosphocreatine, 4 mM ATP-Mg, 0.3 mM GTP-Na, and 0.2 mM EGTA (pH 7.32, 299 mOsm). Neurons were voltage-clamped at −70 mV during membrane rupture, and recordings were excluded if holding currents exceeded −100 pA or the series resistance surpassed 30 MΩ (± 20% fluctuation). Evoked action potentials were characterized by applying depolarizing current steps (−40 to +160 pA, 500 ms) from a −70 mV holding potential.

### Immunohistochemistry

Mice were anesthetized and perfused first with saline and then with 4% paraformaldehyde. The brains were carefully removed, post-perfused with 4% paraformaldehyde for 24 h, and immersed in 30% sucrose solution. A freezing microtome (CM1950; Leica Microsystems) was used to cut 40-μm-thick coronal sections. Free-floating sections were washed in PBS three times, 5 min/wash, permeabilized with 0.3% Triton X-100 in PBS for 30 min, blocked in blocking solution containing 5% normal goat serum and 0.3% Triton X-100 for 2 h at room temperature, and then stained with a rabbit polyclonal anti-TPH2 primary antibody (1:300, NB100-74555; Novus Biologicals) overnight at 4°C. After washing, the sections were incubated with the secondary antibody (1:500, Alexa Fluor 647-conjugated goat anti-rabbit IgG (H+L); Thermo Fisher Scientific) for 2 h, and mounted with a mounting medium containing diamidino-phenylindole (DAPI) (Vectashield; Vector Laboratories, Burlingame, CA, USA).

### Terminal-deoxynucleotidyl transferase-mediated nick end labeling (TUNEL) assay

Apoptosis was detected in brain sections (40 μm thick) using the TUNEL Apoptosis Detection Kit (40308ES50; Yeasen Biotech, Shanghai, China) following the manufacturer’s guidelines. Sections were mounted on glass slides, and washed three times with PBS. Tissue permeabilization was achieved with 0.1% Triton X-100 in PBS (15 min, room temperature), optionally supplemented with Proteinase K (20 μg/mL, 10 min, RT) to enhance DNA accessibility. For positive controls, slides were treated with DNase I (10 U/mL in 1× DNase buffer, 10 min, room temperature) to generate DNA strand breaks. After equilibration in 1× kit-supplied buffer (10 min, room temperature), sections were incubated with a labeling mix containing YSFluor 640-dUTP and TdT enzyme in a humidified chamber (37°C, 60 min, dark) and washed three times with PBS. Nuclei were counterstained with DAPI, and slides were mounted with antifade medium, coverslipped, and sealed with nail polish. Fluorescent imaging was performed using A1R confocal microscope (Nikon Instruments), with specificity confirmed by TdT enzyme omission in negative controls and DNase I-treated sections as positive controls. And TUNEL^+^ cells was quantified using Imaris x64 9.7.1 software (Oxford Instruments).

### Single-cell patch-protein analysis

To detect protein expression in distinct populations of serotonergic projecting neurons at a single-cell level, bilateral stereotaxic injections of AAV11-DIO-EGFP and AAV11-DIO-mCherry were administered into the vCPU and VTA of *Sert*-Cre mice, respectively. Four weeks post-virus injection, a patch-clamp micropipette aspiration technique was used to collect cellular contents from three neuronal subtypes—Type I neurons, mCherry^+^ only, indicating that they project only to the VTA; Type II neurons, EGFP^+^ only, showing that they project only to the vCPU; and Type III neurons, double-positive for mCherry and EGFP, indicating that they doubly project to the VTA and vCPU. Both the solution used for slicing and the ACSF were deoxygenated to simulate hypoxic conditions in the brain and all the solutions were temperature-controlled at 0–4°C. Brain slices were immediately transferred to the recording chamber post-sectioning, with the entire aspiration process completed within 30 min. Electrodes (2–4 MΩ resistance) were filled with 1 μL of an internal solution containing 130 mM K-D-gluconate, 20 mM KCl, 10 mM HEPES, 10 mM phosphocreatine, 4 mM ATP-Mg, 0.3 mM GTP-Na, 0.2 mM EGTA (pH 7.32, 299 mOsm). Aspirated cellular contents were expelled into Eppendorf tubes prefilled with 2 μL of lysis buffer (PMSF: RIPA = 1:100). Two cells were pooled per sample tube. Samples were flash-frozen at –20 or –80°C and promptly processed for Simple Western detection.

### Simple Western assay

For the Simple Western assay, the JESS system (Bio-Techne; ProteinSimple) was used. Samples were prepared by adding 1.5 μL of 5× Mastermix (200 mM dithiothreitol, 5× sample buffer, and fluorescent standards [Standard Pack 1]), denaturing at 95°C for 5 min, and cooling on ice prior to loading. To account for the scarcity of single-cell protein lysates, the loading volume was optimized to 2 μL (*vs*. the standard 3 μL) and the primary antibody incubation time was prolonged from 30 to 60 min. This adjustment to the protocol significantly improved sensitivity for low-abundance targets. For target protein detection, species-matched secondary antibodies were applied (anti-rabbit HRP conjugate [DM-001] or anti-mouse HRP conjugate [DM-002]), with chemiluminescent signals captured in high dynamic range mode. Antibody diluent 2 served as the universal dilution buffer for all immunoreagents. To enable multiplex detection of target proteins and housekeeping proteins in a single capillary, the RePlex stripping protocol was employed. Experimental components—including cellular lysates, antibody diluent 2, primary/secondary antibodies, chemiluminescent substrates (luminol-S/peroxide), RePlex reagent mix, and wash buffer—were transferred to 12–230 kDa prefilled microplates (SM-W004), and then centrifuged at 2500 × *g* for 5 min under ambient conditions. The capillary cartridge and microplate were mounted on their respective holders to initiate automated assays. Compass Software for Simple Western (version 6.3.0, ProteinSimple) was used to operate the Jess system and analyze the results. The separation matrix loading time was set to 200 s, the stacking matrix loading time was set to 15 s, the sample loading time was set to 9 s, the separation time was set to 25 min, the antibody diluent time was set to 5 min, the primary antibody incubation time was set to 60 min, the secondary antibody incubation time was set to 30 min, and the RePlex purge time was set to 30 min. Relative protein concentrations were quantified *via* the peak areas detected in the chemiluminescence electropherogram generated by the Compass for Simple Western software (ProteinSimple). The primary antibodies were used for the analysis were anti-NOX4 (1:100, 14347-1-AP; Proteintech), anti-GPX4 (1:10, ab125066; Abcam), and anti-GAPDH (1:100, 10494-1-AP, Proteintech).

### Behavioral protocols CSDS

CSDS was applied as described in our previous study ^49^. Briefly, experimental mice were physically attacked by different aggressor CD1 mice for 5 min once a day for 10 consecutive days. After each trial, the mouse and the attacker were separated by a perforated translucent plastic partition. Control mice were housed in pairs separated by a partition and were rotated daily, but they were never exposed to attackers. Three distinctive behaviors were defined during exposure to CSDS—attack, approach (the CD1 attacker and the tested mouse stayed within 2 cm of the central partition), and non-approach (the mice remained more than 2 cm from the central partition). Social interaction was assessed 24 h after the last attack. This test was divided into two stages, each lasting 2.5 min. In the first stage, mice were allowed to freely explore a new arena (without a target) in which a small animal cage had been placed on one side. Then, during the second stage, mice were reintroduced into the arena, in which a caged CD1 mouse (with a target) was present. EthoVision XT software (17.0; Noldus, Wageningen, Netherlands) was used to track the length of time the mice stayed in the interaction area. The social interaction (SI) ratio was defined as (interaction time, target) / (interaction time, no target) × 100. Mice with SI scores <100 were categorized as susceptible while those with scores >100 were classified as resilient.

### Restraint stress

Mice were placed in a closed 50-ml tube with ventilation holes for 2 h each day for 21 days. Fiber photometry was performed before and after mice had been placed in the tube on the 1^st^, 7^th^, 14^th^, and 21^st^ day, followed by the forced swimming test (FST), respectively.

### Fear conditioning

Fear conditioning was performed as previously reported ^50^, with minor modifications. Mice were trained and tested using the NIR Video Fear Conditioning System (MED Associated Inc., St. Albans, VT, USA). For training, test cages equipped with stainless steel shock grids connected to the Video Freeze system with precise feedback current regulation. The test cages were placed in a soundproof enclosure (MED Associated Inc.). Behavior was recorded with a low-light camera. Video Freeze software was used to control stimulus presentation. All equipment was thoroughly cleaned with detergent and water after use.

In the fear conditioning paradigm, mice were habituated to the electric shock grid (Context A: metal fence floor, grey metal wall) for 2 min. Fear conditioning was conducted using four pairings of an auditory cue (CS; 30 s, 2800 Hz, 70 dB) co-terminating with a scrambled foot shock (US; 1 s, 0.70 mA). The intervals were randomly set from 60 to 120 s. Freezing duration was recorded during the 30-s CS. After each trial, mice were returned to their home cages. Twenty-four hours later, the contextual test was conducted, in which mice were placed in Context A for 3 min without an auditory cue or foot shock. Freezing behavior was analyzed over 3 min. The cue-dependent fear test was performed 24 h after the contextual test. For this, mice were introduced to a different context (Context B: white Plexiglass floor and walls) for an initial 3-min period, followed by a 3-min presentation of an auditory cue. The mice were returned to their home cage after the end of the tone. The percentage of freezing time during the initial 3 min (BL) and the final 3 min (auditory cue presentation) was calculated.

### SEFL

SEFL was conducted as previously described ^51,52^. Mice were subjected to restraint stress for 7 successive days and then exposed to the training context (MED Associated Inc.) three times in one day (a total of 12 min) for habituation. Twenty-four hours later, mice underwent the following fear conditioning protocol: 2 min of free exploration followed by two 30-s CS–US pairings, separated by a 60-s inter-tone interval, ending with a 0.5-mA foot shock (US). The CS was an 85-dB, 10-kHz pure tone. One minute after the second shock, the mice were returned to their home cages. The freezing behavior within the 1 min after the second shock during fear conditioning training was recorded. Among mice subjected to SEFL, those with above-average freezing percentages were classified as stress-susceptible (SUS), while those with below-average freezing percentages were classified as stress-resilient (RES).

### Mechanical pain stimuli

A 25-G needle was pressed into the center of each hind paw with enough force to indent but not puncture the skin. Each paw was stimulated 10 times, and the HyPer7 signals were recorded before and after stimulation.

### Learning helplessness

The method used for the learning-helplessness paradigm was as previously reported ^53^. Mice were trained for 3 days (2 h each session) on one side of a two-chamber shuttle box (MED Associated, Inc.), during which time the door between the chambers was closed and the animals were given 120 shocks at different internals (5–99 s; 0.3 mA for 5 s). On the day of the test, the door was raised at the start of the shock, and the trial ended when the mice walked to the other side of the shuttle box or after 25 s. The latency to step through the door and the number of escape failures were recorded for 25 trials.

### FST

Mice were gently placed in a transparent glass cylinder (height 45 cm, diameter 19 cm) filled with water (22–25°C) to a depth of 23 cm. The immobility time was recorded during the last 4 min of a 6-min test session using EthoVision XT software.

### Open field test (OFT)

Mice were placed in the central zone of the SuperFlex Open Field System (Omnitech Electronics, Inc., Columbus, OH, USA) and were permitted to freely explore for 5 min. The total distance traveled and time spent in the center of the arena were analyzed using Fusion software (6.5; Omnitech Electronics, Inc.).

### Light–dark box (LDB) test

For the light–dark box test, the same chamber as that used for the OFT was divided into two equally sized compartments, one bright and one dark, with a hole in the partition wall that allowed free passage between the compartments. The mice were placed in the center of the bright chamber with their backs to the hole and allowed to freely explore for 5 min. Fusion software was used to analyze the time spent in the dark compartment and the number of transitions between the two compartments.

### Elevated plus maze (EPM) test

The elevated plus maze consisted of two opposite open arms (30 × 5 × 0.5 cm) and two opposite closed arms (30 × 5 × 15 cm) connected by a central platform (5.5 × 5.5 cm). The mice were gently placed in the center of the maze and allowed to explore for 5 min. The time spent in the open arms and the number of entries into the open arms were quantified using EthoVision XT software.

### Sucrose preference test (SPT)

Mice were acclimated for 4 days to two 100 mL bottles fitted with ballpoint sipper tubes. The bottles were filled with drinking water in the first 2 days and with 1% sucrose (V900116; Sigma-Aldrich) for the next 2 days. After habituation, the mice were provided with one bottle containing 1% sucrose and one containing only water. Sucrose preference was calculated as the percentage of sucrose solution consumed relative to the total liquid intake using the formula (amount of sucrose consumed × 100 [bottle A] / total volume consumed [bottles A + B]).

### Novelty-suppressed feeding test (NSFT)

After fasting for 24 h, mice were placed in a plastic box (50 × 50 × 20 cm) containing 2-cm-thick wooden bedding. The NSFT lasted for 5 min. At the beginning of each test, a food particle was placed on a white paper platform in the center of the box. Mice were gently placed in a corner of the box, and the time it took the mouse to reach for the food with its forepaws and start eating was recorded. After the experiment, the mice were immediately placed in their home cages, and their food consumption was measured for the next 10 min.

### Y-maze test

The Y-maze was made of gray Plexiglass and had three identical 30 × 10 × 20 cm arms and a central triangular area measuring 10 × 10 × 10 cm connecting the three arms. Each arm formed a 120° angle with the other two arms. During a trial, mice were placed in the central area and allowed to freely explore the maze for 5 min. The sequence of arm entries was recorded. An alternation was defined as three consecutive entries into different arms and was scored as one point. The total number of opportunities for alternation was calculated as the total number of arm entries minus two. The percentage of alternations was then determined using the following formula: (total alternation points / total alternation opportunities) × 100.

## QUANTIFICATION AND STATISTICAL ANALYSIS

### Calculation of O_2_ utilization

The proportion of O_2_ utilization was calculated in the three types of DRN^5-HT^ neurons after the CSDS paradigm based on NOX4 expression and cell counts. The proportion of NOX4 among three type neurons was calculated by the results obtained from the single-cell patch-protein analysis multiplied by the number of cells survived in each respective neuron type, yielding the total expression level of NOX4 in each type of neuron. The proportion of O_2_ utilization in each of the three types of neurons was calculated as percentages for each behavioral phenotype.

To assess the effects of the overexpression of *NOX4* in type III neurons, the O_2_ utilization pattern was evaluated. The total O_2_ utilization in each type neuron was determined by multiplying the DHE fluorescence intensity of a single cell by the number of cells survived. The proportion of O_2_ utilization in each cell type was then calculated as a percentage for each behavioral phenotype.

### Statistical analysis

All data were analyzed using GraphPad Prism (10.0; GraphPad Software, San Diego, CA, USA). Data are presented as means ± SEMs (details are provided in **Supplementary Table 4**). Animals with missed virus injection, incorrect optical fiber placement, or incorrect cannula placement were excluded from analysis by experimenters blinded to the treatment groups. Two-tailed unpaired *t*-tests or multiple *t*-tests were used for comparisons between two groups. Three or more groups were compared by one-way ANOVA, followed by Tukey’s post hoc multiple comparisons test. Two-way ANOVA followed by Bonferroni’s multiple-comparisons test, Sidak’s multiple comparisons test or Tukey’s multiple comparisons test were used for comparisons involving two factors. Two-way repeated-measure ANOVA, followed by Sidak’s multiple comparisons test, was used for repeated measures (RM) analysis. One sample *t*-tests were used to calculate the rHF z-score relative to the basal level. Data distribution was assumed to be normal when ANOVA and *t*-tests were performed. Differences were considered significant if *p* or *q* < 0.05.

## Supporting information

Table S1-S3. Biological samples

Table S4. Summary of statistical analyses

S5. Original WB

## ACKNOWLEDGMENTS

We thank Qian-wen Zhao, Xue-Fang Zeng and Hao Liu, Pazhou Lab, for their technical supports. This work was supported by the STI 2030 – Major project 2022ZD0211700 (X.-H.Z.), the National Natural Science Foundation of China 82530037 (X.-H.Z.), the Outstanding Talents of Yangcheng Elite Plan 2024D02J0003 (X.-H.Z.), the Guangdong Tezhi plan talent 2024JC08Y144 (X.-H.Z.), and the Guangdong Natural Science Foundation of China 2025A1515012292 (Y.Z.).

## AUTHOR CONTRIBUTIONS

X.-H.Z. conceived and supervised the project. X.-H.Z., Y.Z., and Y.-D.P. designed the experiments and wrote the paper. Y.Z., W.-Y.Z., and Q.-Q.W. carried out the DHE staining. Y.Z. and Y.-D.P. performed experiments for stereotaxic microinjections and optical fiber/cannula implantation. Y.Z. and Y.-D.P. performed NADH/NAD+ analysis and 5-HT and 5-HIAA detection. Y.Z., W.-Y.Z., and M.-Z.Z. performed experiments for HyPer7 detection. Y.Z., Y.-D.P., and Q.-Q.W. performed PCR, western blot analysis, prepared brain slices and immunostaining. Y.-D.P. performed the in vitro electrophysiological recording. Y.Z. and Y.-D.P. constructed the single cell patch-protein analysis. Y.Z., Y.-D.P., W.-Y.Z., Q.-Q.W., H.-Y.L., X.-R.Y., Y.W., and J.-H.D. conducted the behavioral tests. Y.Z., Y.-D.P., W.-Q.L., Z.-Y.Z., and B.-C.L. bred the mice and performed the genotyping. G.T. and N. M. characterized and prepared the postmortem brain samples, and M.D. detected NOX4 protein levels.

