## Supplementary information for "Serotonergic neurons in the dorsal raphe nucleus scale O₂ utilization in response to traumatic stress"

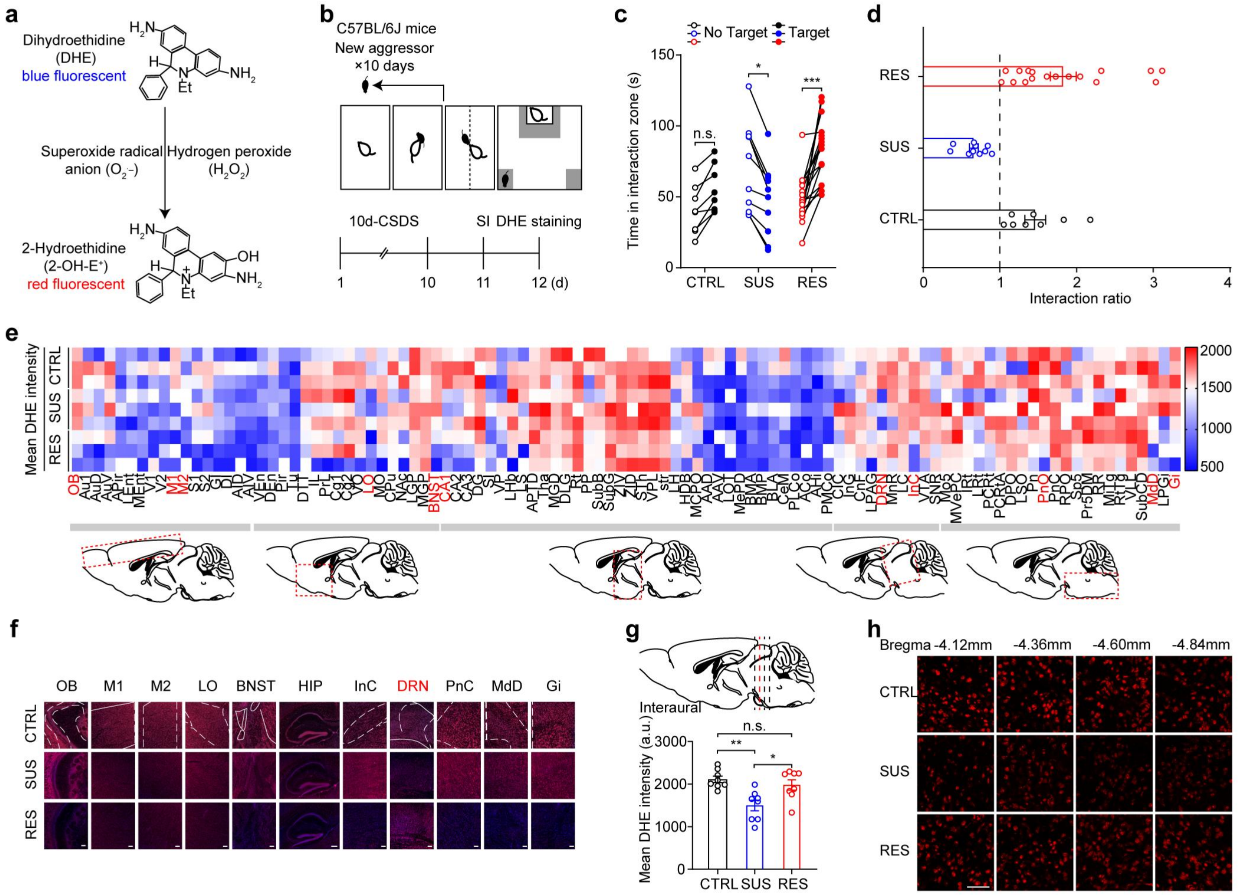


**Supplementary Fig. 1: DHE staining in the brain of C57BL/6J mice following a 10-day CSDS paradigm.**

**a.** Working model of dihydroethidium (DHE). ROS react with DHE to produce 2-hydroxyethidium.

**b.** Schematic of the social interaction (SI) test after a 10-day CSDS paradigm. DHE staining was conducted 24 h after the SI test.

**c.** Social time in the interaction zone after a 10-day CSDS paradigm ($n_{CTRL}$ = 8; $n_{SUS}$ = 10; $n_{RES}$ = 17).

**d.** SI ratio in CSDS ($n_{CTRL}$ = 8; $n_{SUS}$ = 10; $n_{RES}$ = 17).

**e.** Quantification of DHE fluorescence intensity in whole brain areas ($n$ = 3 for each phenotype). One-way ANOVA with Tukey's multiple comparisons test was used for each brain area. Abbreviations for all brain regions refer to Paxinos & Franklin, The Mouse Brain in Stereotaxic Coordinates, Second Edition (2001).

**f.** Representative fluorescence images of oxidized DHE in brain regions showing significant differences in (**e**) (red color) in SUS, RES, and CTRL animals.

**g, h.** DHE fluorescence images and quantification of DHE intensity using coronal brain slice analysis (*n* = 8). The dashed lines indicate bregma levels, and the red line indicates that representative images were collected at bregma –4.36 mm throughout the main text.

Scale bars, 100 μm; $^{*}p < 0.05$, $^{**}p < 0.01$, and $^{***}p < 0.001$.

**c,** two-way ANOVA with Bonferroni's multiple comparisons test. **e, g,** one-way ANOVA with Tukey's multiple comparisons test. n.s., not significant. Data are presented as mean ± SEM. Statistical analysis parameters are included in Supplementary Table 4 hereafter.

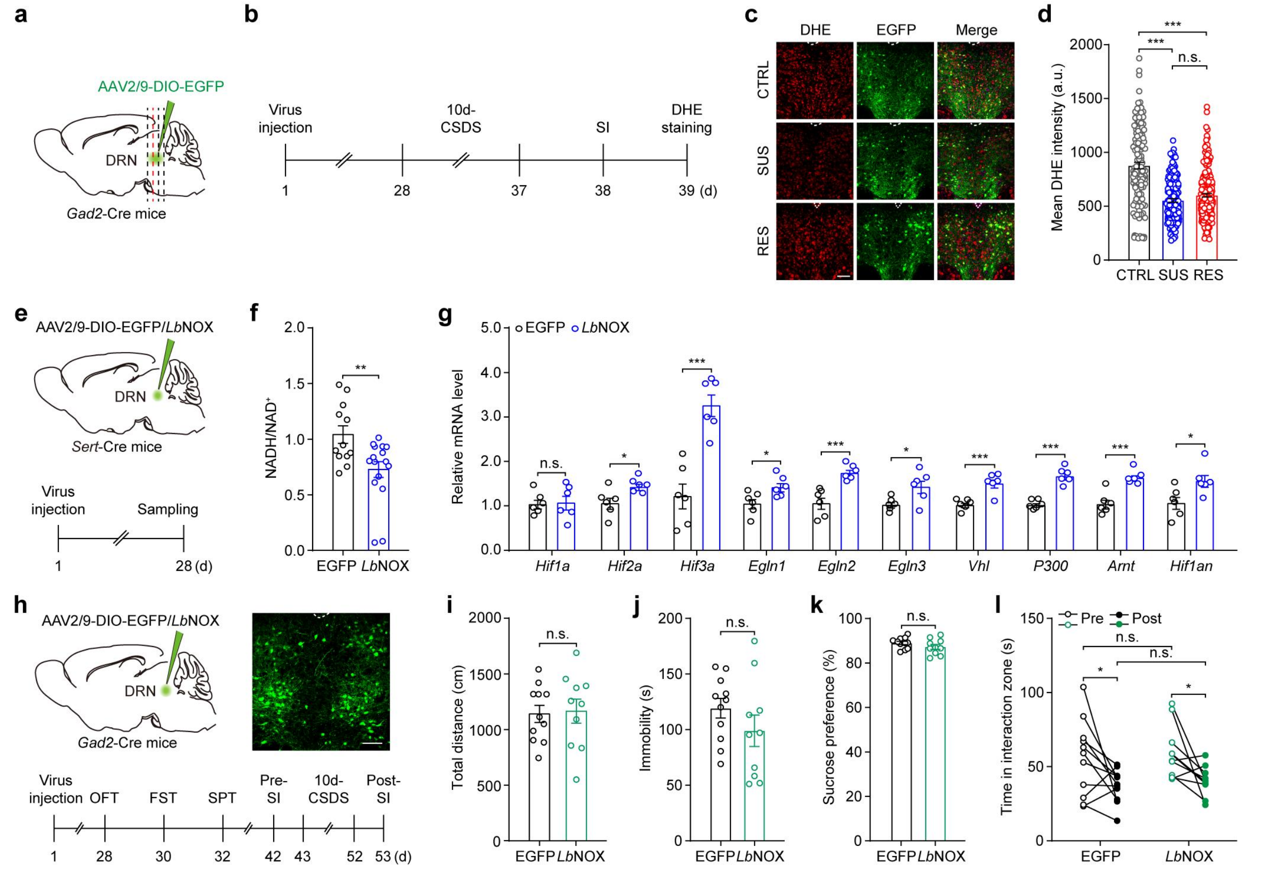


**Supplementary Fig. 2: Expression of *Lb*NOX inhibits activation of the HOSP in DRN$^{5\text{-}HT}$ neurons.**

**a, b.** Schematic of virus injection into the DRN of *Gad2*-Cre mice.

**c, d.** Oxidized DHE fluorescence in DRN$^{GABA}$ neurons after 10 days of CSDS ($n$ = 6 mice per group).

**e.** Schematic of virus injection. AAV2/9-DIO-*Lb*NOX (*Lb*NOX) or control virus (AAV2/9-DIO-EGFP) was injected into the DRN of *Sert*-Cre mice.

**f.** Effects of *Lb*NOX expression on the NADH/NAD$^+$ ratio in the DRN ($n_{EGFP}$ = 12; $n_{LbNOX}$ = 16).

**g.** Expression of genes related to the HOSP in the DRN following *Lb*NOX expression ($n$ = 6).

**h–l.** Experimental timeline of AAV2/9-DIO-*Lb*NOX or AAV2/9-DIO-EGFP injection into the DRN of *Gad2*-Cre mice and subsequent behavioral tests (**h**) ($n_{EGFP}$ = 11, $n_{LbNOX}$ = 10).

Total distance traveled in the OFT (**i**), immobility in the FST (**j**), SPT ratio (**k**), and social interaction test before (Pre) or after (Post) a 10-day CSDS (**l**).

Scale bars, 100 μm; $^{*}p < 0.05$, $^{**}p < 0.01$, and $^{***}p < 0.001$.

**d,** one-way ANOVA with Tukey's multiple comparisons test. **f, i, j, k,** two-tailed unpaired *t*-test. **g,** multiple *t*-test. **l,** two-way ANOVA with Sidak's multiple comparisons test. n.s., not significant. Data are presented as mean ± SEM.

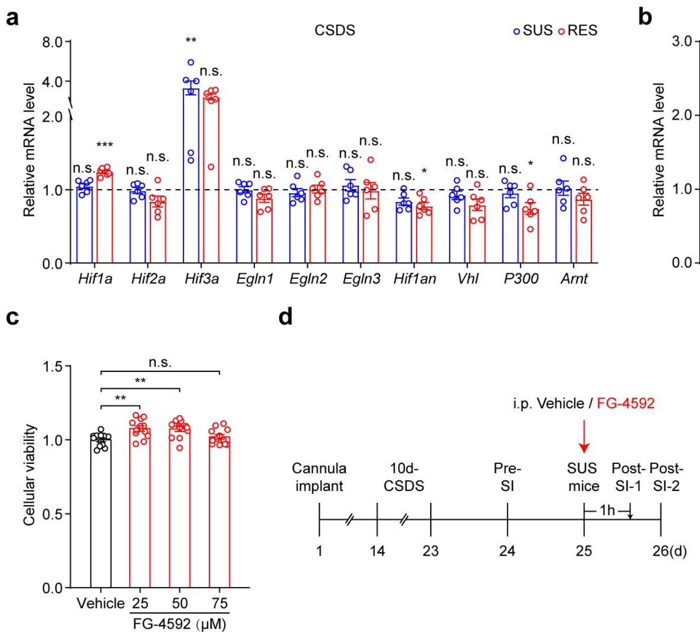


**Supplementary Fig. 3: Infusion of low-dose FG-4592 into the DRN has little effect on depressive-like behaviors in CSDS paradigm.**

**a, b.** qPCR analysis of HOSP-related gene expression in the DRN after a 10-day CSDS paradigm (**a**) ($n$ = 6) and SEFL paradigm (**b**) ($n$ = 6). The mRNA level of *Hif3a* is too low in the DRN to be analyzed further, as its *Ct* value is approximately equal to 32.33.

**c.** Effects of different concentrations of FG-4592 on the viability of cultured HEK293 cells, as detected via a CCK–8 assay ($n$ = 2 technical replicates from 6 biological replicates for each dosage).

**d–f.** Experimental timeline for DRN-infusion of FG-4592 (2.5 μM) in CSDS. After cannula implantation, C57BL/6J mice were subjected to a 10-day CSDS paradigm, and SUS mice were given only one injection of FG-4592, and behaviors were detected 1 h or 24 h after treatment. Time in the social interaction zone 1 h (SI-1) (**e**) and 24 h (SI-2) (**f**) after the infusion of FG-4592 ($n$ = 8).

$^{*}p < 0.05$, $^{**}p < 0.01$, and $^{***}p < 0.001$.

**a, b, c,** one-way ANOVA with Tukey's multiple comparisons test. **e, f,** two-way ANOVA with Sidak’s multiple-comparisons test. n.s., not significant. Data are presented as mean ± SEM.

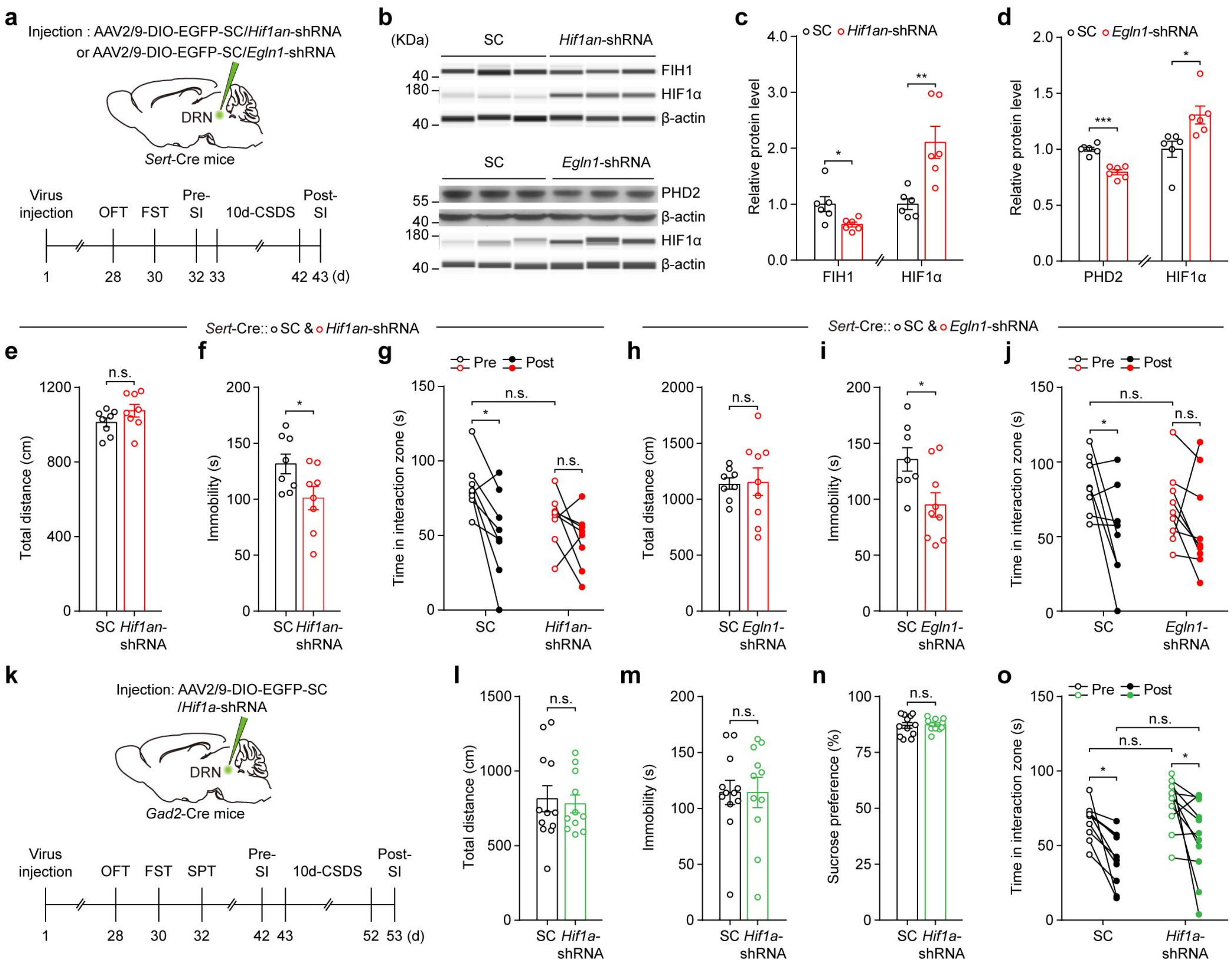


**Supplementary Fig. 4: Behavioral effects of viral modulation of the HOSP in DRN neurons.**

**a.** Schematic of viral injection for expressing *Hif1a*-shRNA or *Egln1*-shRNA in DRN$^{5\text{-HT}}$ neurons. Control virus, AAV2/9-DIO-EGFP–scrambled sequence (SC).

**b–d.** Simple western blot images (**b**) and quantification of the levels of FIH1 (**c**), PHD2 (**d**), and HIF1α in the DRN 28 days after virus injection ($n$ = 6).

**e–j.** Behavioral tests for knocking down FIH1 (**e–g**) ($n$ = 8) and PHD2 (**h–j**) ($n_{\text{SC}}$ = 8; $n_{Egln1\text{-shRNA}}$ = 9) in DRN$^{5\text{-HT}}$ neurons, conducted 28 days after virus injection. Effect of HIF1α upregulation in DRN$^{5\text{-HT}}$ neurons on locomotion in the OFT (**e, h**), total immobility time in the FST (**f, i**), and social interaction test before (Pre) or after (Post) a 10-day

CSDS paradigm (**g, j**).

**k–o.** Schematic of *Hif1a*-shRNA expression in DRN$^{GABA}$ neurons. AAV2/9-DIO-EGFP–*Hif1a*-shRNA or control virus was injected into the DRN of *Gad2*-Cre mice, and behaviors were evaluated 28 days after virus injection (**k**) ($n_{SC}$ = 10–12; $n_{Hif1a\text{-shRNA}}$ = 11). Total distance traveled in the OFT (**l**), immobility in the FST (**m**), SPT ratio (**n**), and social interaction test before (Pre) and after (Post) a 10-day CSDS paradigm (**o**).

$^{*}p < 0.05$, $^{**}p < 0.01$, and $^{***}p < 0.001$.

**c, d, e, f, h, i, l, m, n,** two-tailed unpaired *t*-test. **g, j, o,** two-way ANOVA with Sidak's multiple comparisons test. n.s., not significant. Data are presented as mean ± SEM.

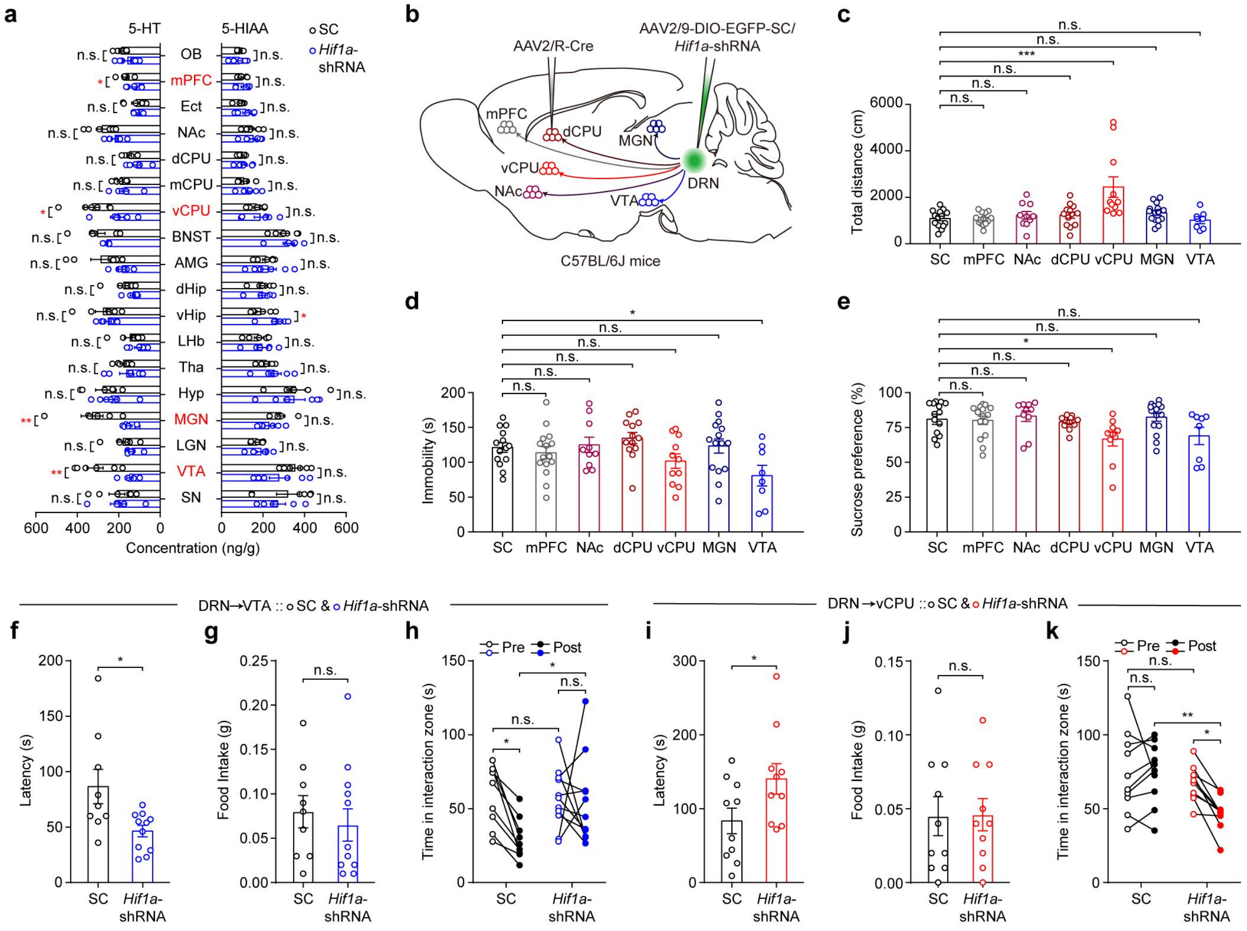


**Supplementary Fig. 5: Knocking down *Hif1a* in DRN neurons projecting to the vCPU or VTA in C57BL/6J mice differentially modulates depression-related behaviors.**

**a.** 5-HT and 5-hydroxyindoleacetic acid (5-HIAA) concentrations detected in various brain regions after knocking down *Hif1a* in DRN$^{5\text{-HT}}$ neurons of *Sert*-Cre mice ($n$ = 6). AAV2/9-DIO-EGFP–*Hif1a*-shRNA or control virus was injected into the DRN, and concentrations of 5-HT and its metabolite were detected using HPLC-ECD analysis 28 days after virus injection.

**b–e.** Schematic of *Hif1a*-shRNA manipulation of DRN neurons projecting to the mPFC, NAc, vCPU (ventral part of CPU), dCPU (dorsal part of CPU), VTA, and MGN, and behavioral tests. AAV2/9-DIO-EGFP–SC or AAV2/9-DIO-EGFP–*Hif1a*-shRNA was injected into the DRN of C57BL/6J mice, and AAV2/R-Cre virus was injected into the aforementioned brain areas (**b**). Behavioral tests were conducted 28 days after virus injection ($n_{SC}$ = 14; $n_{mPFC}$ = 15; $n_{NAc}$ = 10; $n_{dCPU}$ = 13; $n_{vCPU}$ = 11; $n_{MGN}$ = 15; $n_{VTA}$ = 8). Total distance traveled in the OFT (**c**), immobility in the FST (**d**), and SPT ratio (**e**).

**f–h.** Behavioral effects of *Hif1a* knockdown in DRN→VTA neurons ($n_{SC}$ = 9; $n_{Hif1a\text{-shRNA}}$ = 11): latency in the NSFT (**f**), food consumption (**g**), and social interaction tests after a 10-day CSDS paradigm (**h**).

**i–k.** Effects of downregulation of HIF1α in DRN→vCPU neurons (*n* = 10): latency in the NSFT (**i**), food intake (**j**), and social interaction tests after a 3-day CSDS paradigm (**k**). $^{*}p < 0.05$, $^{**}p < 0.01$, and $^{***}p < 0.001$.

**c, d, e,** one-way ANOVA with Tukey’s multiple comparisons test. **f, g, i, j,** two-tailed unpaired *t*-test. **h, k,** two-way ANOVA with Sidak’s multiple comparisons test. n.s., not significant. Data are presented as mean ± SEM.

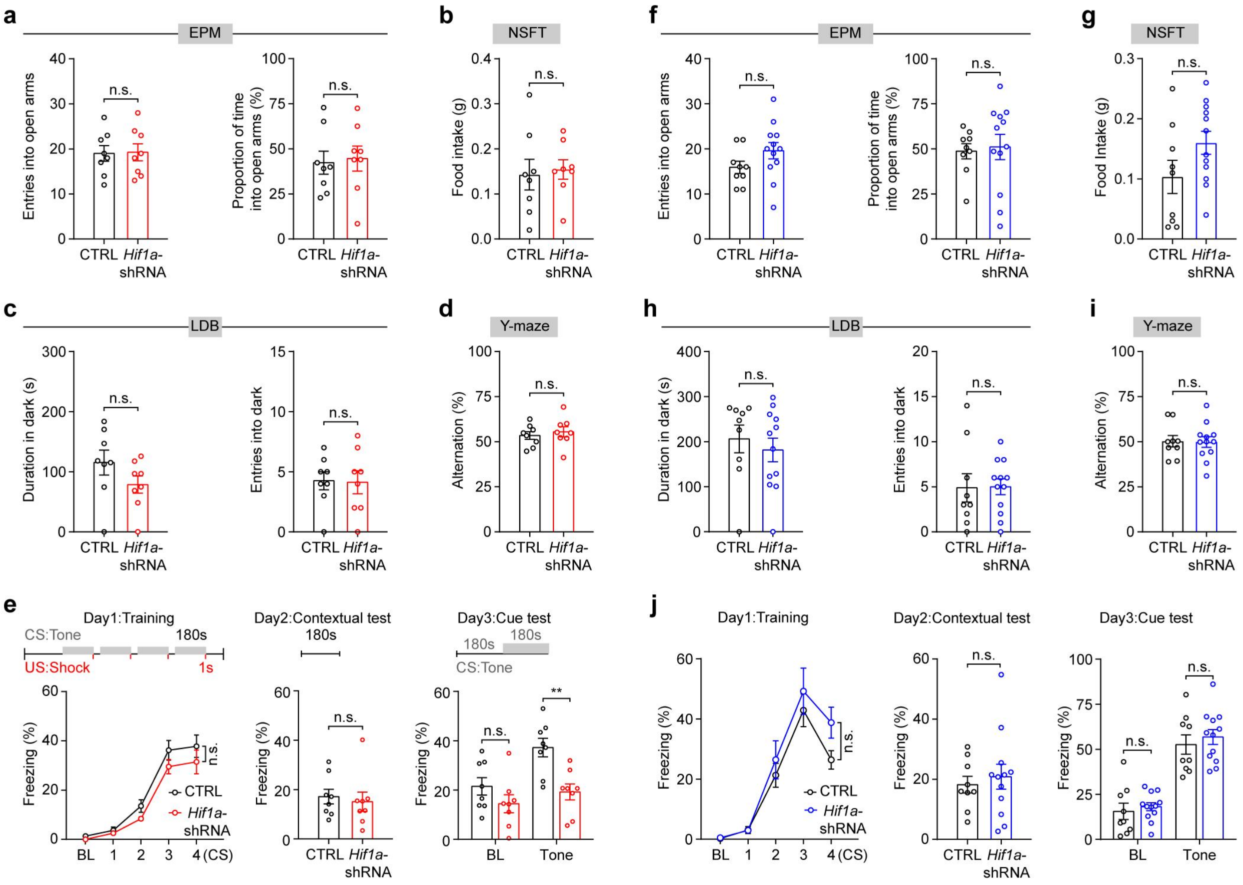


**Supplementary Fig. 6: Expression of *Hif1a*-shRNA in Sert$^{DRN \rightarrow vCPU}$ or Sert$^{DRN \rightarrow VTA}$ neurons has little effect on anxiety-like behaviors and cognition.**

**a–j.** Behavioral effects of HIF1α downregulation in Sert$^{DRN \rightarrow vCPU}$ neurons 28 days after virus injection (**a–e**) ($n$ = 8) and behavioral effects of *Hif1a* knockdown in Sert$^{DRN \rightarrow VTA}$ neurons (**f–j**) ($n_{SC}$ = 9; $n_{Hif1a\text{-shRNA}}$ = 12).

Number of entries into the open arms (left) and percentage of time spent in the open arms (right) in the elevated plus maze (EPM) test (**a, f**).

Food intake in the NSFT (**b, g**).

Duration in the dark chamber in the light–dark box (LDB) test (**c, h**).

Alternation in the Y-maze test (**d, i**).

Behavioral effects of *Hif1a* downregulation in the fear conditioning test (**e, j**).

Downregulation of HIF1α in Sert$^{DRN \rightarrow vCPU}$ neurons decreased freezing time in the cue test (**e**), possibly due to hyperactivity. Downregulation of HIF1α in Sert$^{DRN \rightarrow VTA}$ neurons had no effect on freezing behaviors during fear training or contextual and cue tests (**j**). $^{**}p < 0.01$.

**a, b, c, d, f, g, h, i,** two-tailed unpaired *t*-test and the contextual and cue tests in (**e, j**). **e, j,** two-way RM ANOVA with Sidak's multiple comparisons test. n.s., not significant. Data are presented as mean ± SEM.

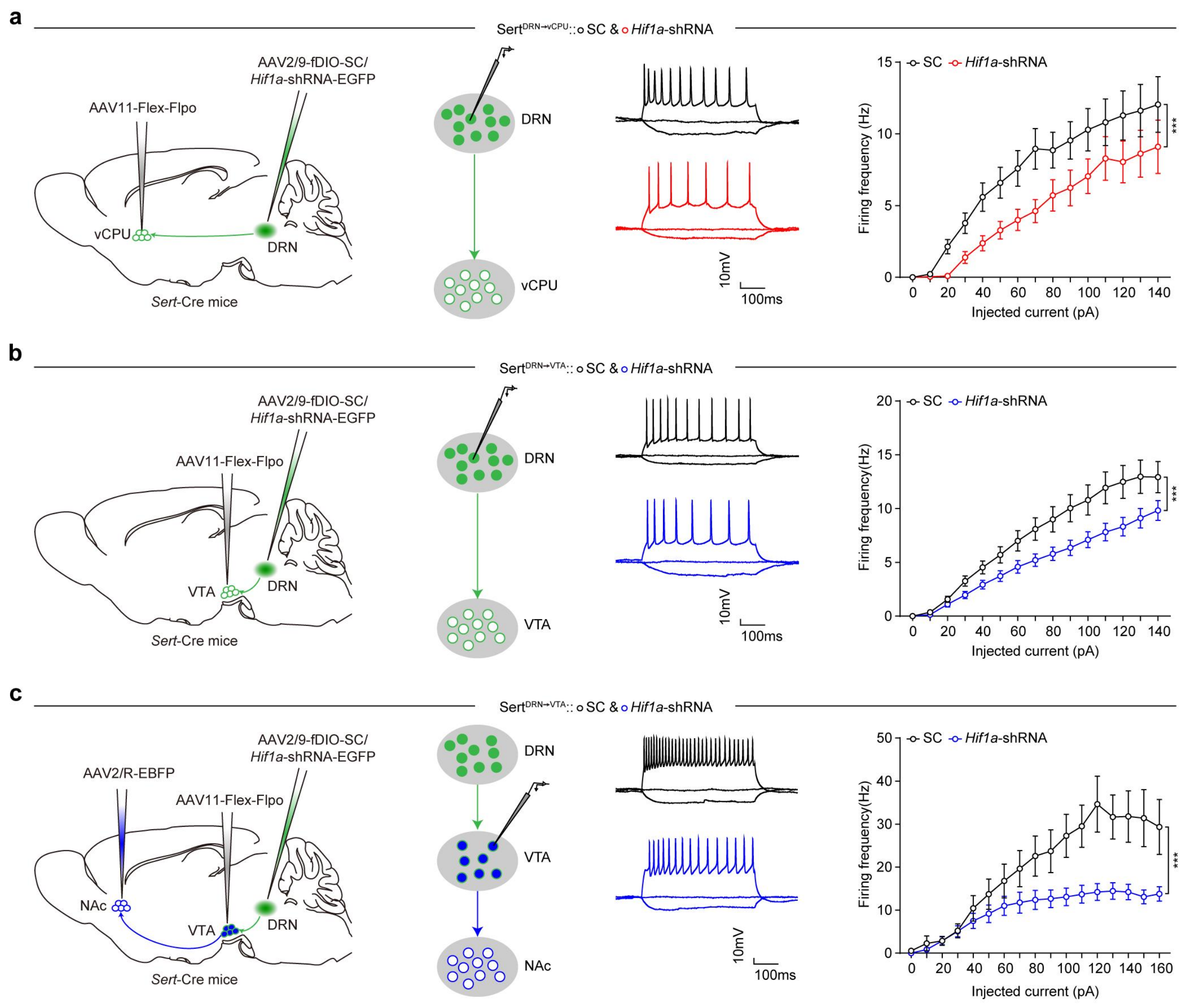


**Supplementary Fig. 7: The HOSP regulates activation of DRN$^{5\text{-HT}}$ neurons.**

**a, b.** Whole-cell electrophysiological recordings showed that knocking down *Hif1a* in Sert$^{DRN\rightarrow vCPU}$ (**a**) ($n$ = 21–22 cells from three mice per group) and Sert$^{DRN\rightarrow VTA}$ (**b**) ($n$ = 29–30 cells from three mice per group) neurons decreased their firing rates, respectively.

**c,** Downregulation of HIF1α in Sert$^{DRN\rightarrow VTA}$ neurons decreased the firing rates of VTA neurons projecting to the NAc ($n$ = 15–24 cells from three mice per group).

$^{***}p < 0.001$.

**a, b, c,** two-way ANOVA with Bonferroni's multiple comparisons test. Data are presented as mean ± SEM.

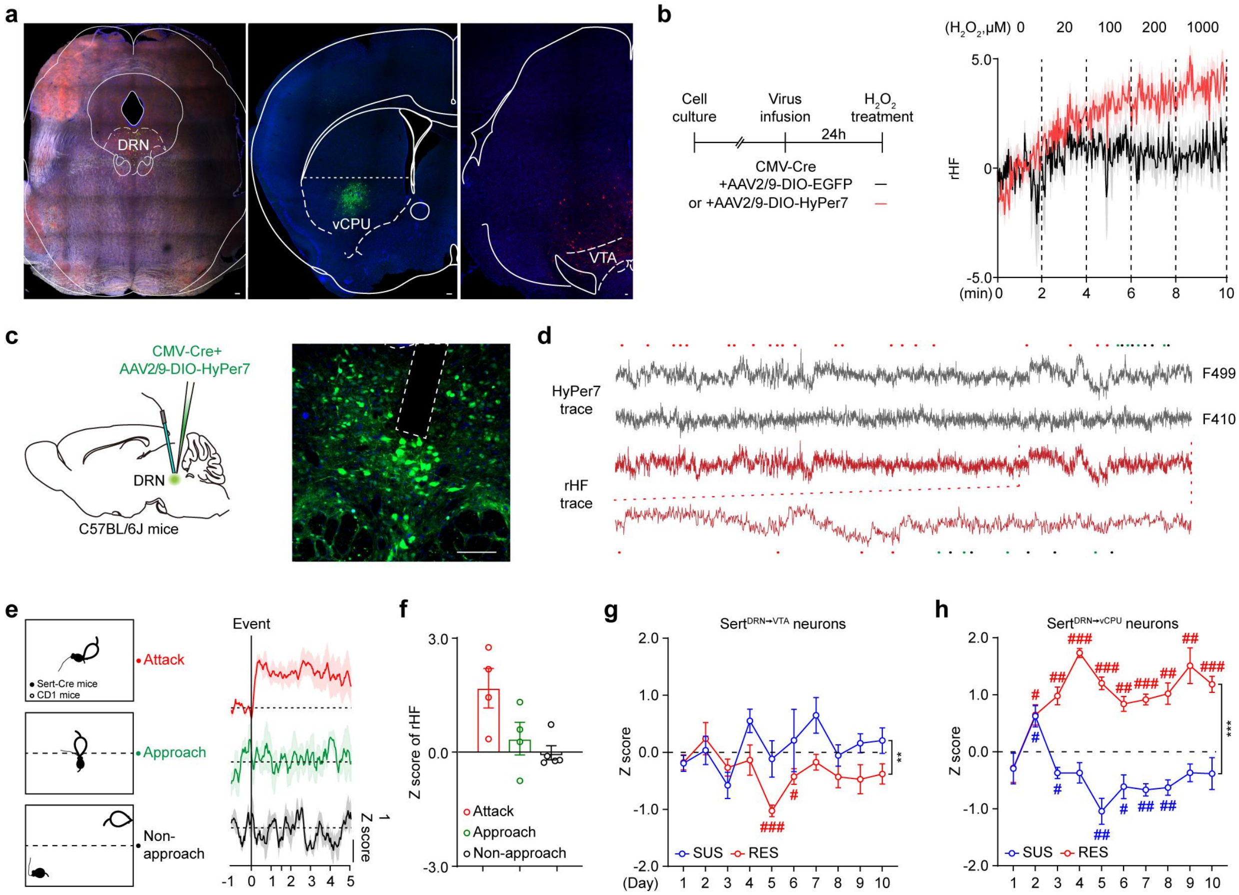


**Supplementary Fig. 8: HyPer7 fluorescence intensity recordings.**

**a.** Confocal images of labeled Sert$^{DRN \rightarrow VTA}$ (red) and Sert$^{DRN \rightarrow vCPU}$ (green) neurons. The cell bodies of these neurons were gathered in the DRN, and terminal fibers were observed in the vCPU (EGFP) or VTA (mCherry), respectively.

**b.** Time course of changes in the ratio of HyPer7 fluorescence intensity (rHF) upon the sequential addition of $H_2O_2$ at concentrations of 20, 100, 200, or 1000 μM ($n$ = 3 technical replicates from six biological replicates for each dosage).

**c, d.** Example data extracted from digitally captured HyPer7 fluorescence signals in the DRN of adult C57BL/6J mice following CSDS. Changes in fluorescence signal over time (**d**), represented as fluorescence at each time point; the zoomed portion shows example rHF corresponding to three distinct behaviors in one mouse. Red dots, attacks; green dots, approaches; black dots, non-approaches.

**e.** Images showing rHF traces in response to three distinctive behaviors following CSDS. Left, diagrams of attack, approach, and non-approach behaviors in CSDS. Right, representative traces of rHF.

**f.** Calculated mean z-score of rHF in response to three distinctive behaviors.

**g, h.** Quantification of mean attack-locked rHF in $Sert^{DRN \rightarrow VTA}$ (*n* = 5/8 for SUS/RES mice) (**g**) and $Sert^{DRN \rightarrow vCPU}$ (*n* = 6/6 for type II neurons in SUS/RES mice) (**h**) during attack bouts throughout a 10-day CSDS.

Scale bars, 100 μm; $^{**}p < 0.01$ and $^{***}p < 0.001$.

**g, h,** two-way RM ANOVA with Sidak's multiple comparisons test. $^{\#}p < 0.05$, $^{\#\#}p < 0.01$, and $^{\#\#\#}p < 0.001$ relative to the basal level (dashed line) by one-sample *t*-test (**g, h**).

Data are presented as mean ± SEM.

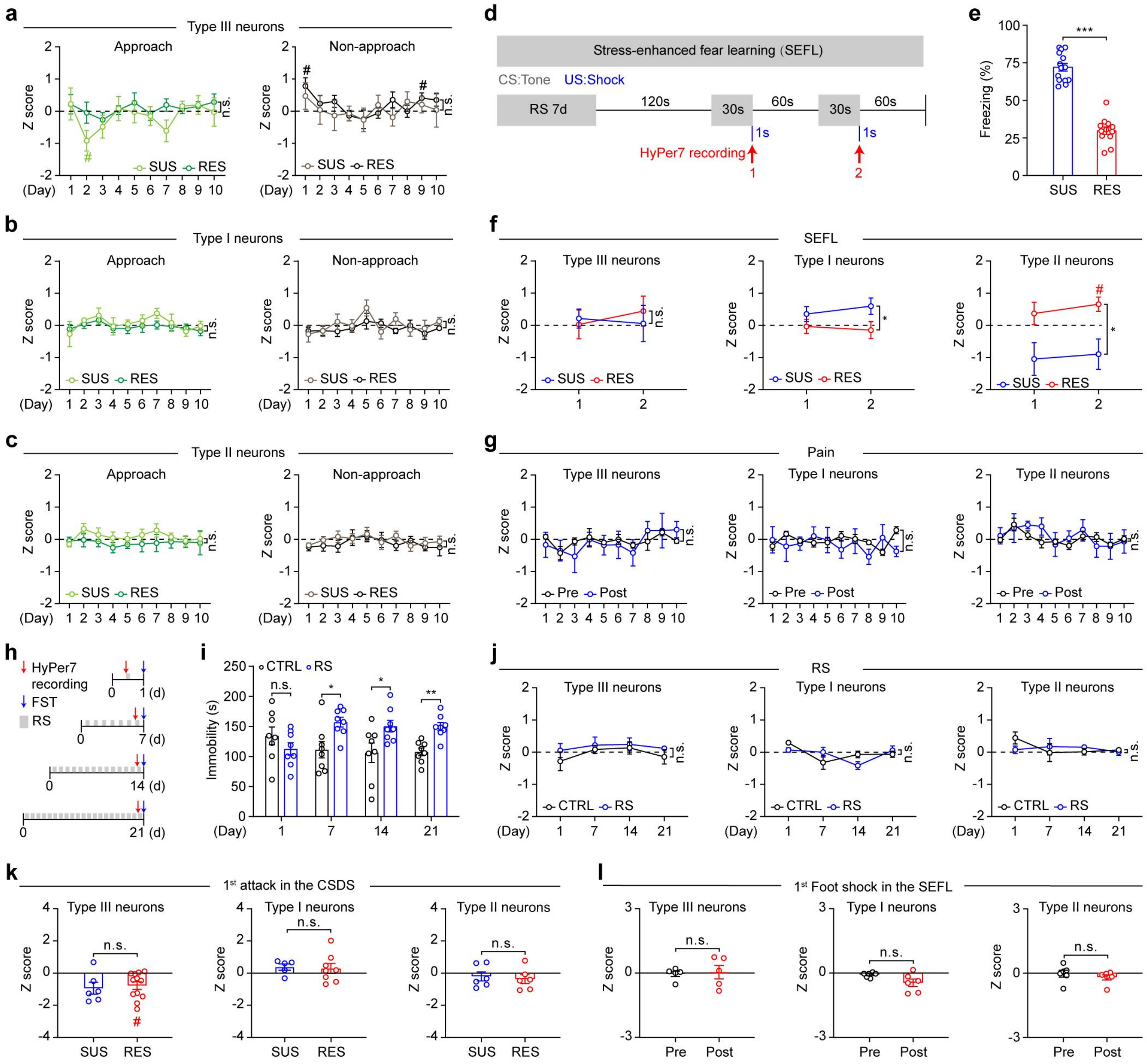


**Supplementary Fig. 9: The rHF of DRN$^{5\text{-HT}}$ neurons does not change in response to chronic restraint stress or pain stimuli.**

**a–c.** Quantification of mean event-locked rHF in three types of DRN$^{5\text{-HT}}$ neurons during approach and non-approach bouts in a 10-day CSDS paradigm (*n* = 6/12 for type III neurons in SUS/RES mice; *n* = 5/8 for type I neurons in SUS/RES mice; *n* = 6/6 for type II neurons in SUS/RES mice).

**d.** HyPer7 recording during the SEFL paradigm. Three types of DRN$^{5\text{-HT}}$ neurons were labeled following the procedures in Fig. 6d, 6g and 6j. Four weeks after virus injection, *Sert*-Cre mice were subjected to restraint stress (RS) once a day for seven consecutive days, and then given a tone paired with foot shock twice. HyPer7 fluorescence intensity was recorded during tone and foot shock, and the mean z-score of rHF for each type of $DRN^{5\text{-}HT}$ neuron was calculated.

**e.** Behavioral test after the SEFL paradigm ($n$ = 12–14).

**f.** Mean rHF z-score in SUS and RES mice following the SEFL paradigm ($n$ = 5/5 for type III neurons in SUS/RES mice; $n$ = 7/6 for type I neurons in SUS/RES mice; $n$ = 6/5 for type II neurons in SUS/RES mice).

**g.** Quantification of mean event-locked rHF during pain stimulation ($n$ = 5 for type III neurons in mice; $n$ = 7 for type I neurons in mice; $n$ = 7 for type II neurons in mice).

**h.** Schematic for HyPer7 recording in the RS paradigm.

**i.** Changes in immobility time in the FST following RS ($n$ = 8).

**j.** Fiber photometry recordings of HyPer7 following RS ($n$ = 8/8 for type III neurons in CTRL/RS mice; $n$ = 6/7 for type I neurons in CTRL/RS mice; $n$ = 6/7 for type II neurons in CTRL/RS mice).

**k, l.** HyPer7 recordings in three types of $DRN^{5\text{-}HT}$ neurons in response to the first attack (**k**) in the CSDS paradigm ($n$ = 6/12 for type III neurons in SUS/RES mice; $n$ = 5/8 for type I neurons; $n$ = 6/6 for type II neurons) and the first foot shock (**l**) in the SEFL paradigm ($n$ = 5/5 for type III neurons; $n$ = 6/6 for type I; $n$ = 6/6 for type II).

$^{*}p < 0.05$, $^{**}p < 0.01$, $**p < 0.001$.

**a, b, c, f, g, j,** two-way RM ANOVA with Sidak's multiple-comparisons test. **e, k, l,** two tailed unpaired *t*-test. **i,** multiple unpaired *t*-test. n.s., not significant. $^{\#}p < 0.05$ relative to the basal level by one sample *t*-test (**a, f, k**). Data are presented as mean ± SEM.

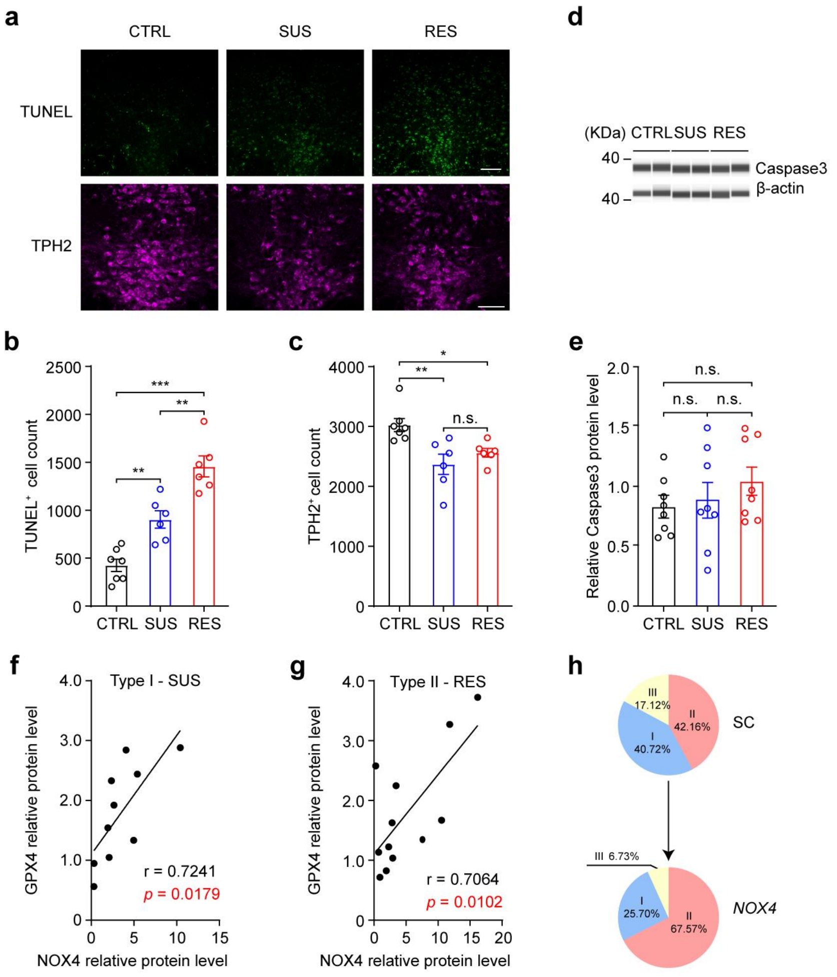


**Supplementary Fig. 10: Cell death in the DRN following the 10-day CSDS paradigm.**

**a–c.** Representative images (**a**) and quantification of the numbers of TUNEL⁺ cells (**b**) and TPH2⁺ cells (**c**) in the DRN of C57BL/6J mice following a 10-day CSDS paradigm ($n_{CTRL}$ = 7; $n_{SUS}$ = 6; $n_{RES}$ = 6).

**d, e.** Simple western blot analysis of Caspase-3 expression in the DRN of C57BL/6J mice following a 10-day CSDS paradigm and control mice (*n* = 8).

**f, g.** Correlations between NOX4 expression and GPX4 protein levels in type I neurons of SUS mice (**f**) or type II neurons of RES mice (**g**) following a 10-day CSDS paradigm.

Pearson's correlation test.

**h,** Overexpressing *NOX4* in type III neurons induced a RES-like $O_2$ scaling pattern in the DRN. The proportion of oxygen utilization for each type of $DRN^{5\text{-}HT}$ neuron was calculated by multiplying the number of surviving neurons by the DHE intensity in each cell, then dividing by the total oxygen utilization of all these neurons [(surviving neurons × DHE intensity per cell) / total oxygen utilization × 100].

Scale bars, 100 μm; $^{*}p < 0.05$, $^{**}p < 0.01$, $^{**}p < 0.001$.

**b, c, e,** one-way ANOVA with Tukey's multiple comparisons test. n.s., not significant.

Data are presented as mean ± SEM.

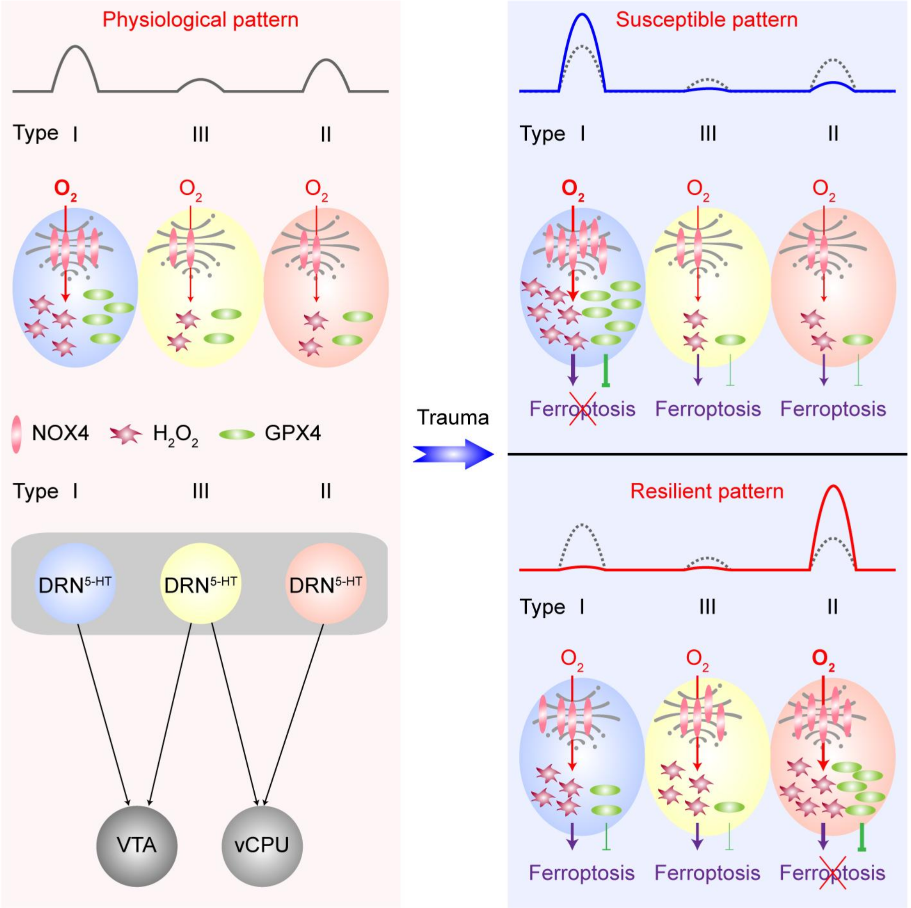


**Supplementary Fig. 11: A working model illustrating a novel function of $O_2$ in the brain.**

Serotonergic neurons in the dorsal raphe nucleus projecting to the ventral caudate putamen and the ventral tegmental area exhibit distinct $O_2$ utilization patterns under physiological conditions and scale $O_2$ utilization in response to traumatic stress following either the susceptible or resilient pattern, thereby modulating corresponding behavioral outcomes.
