## Supplementary material for "Serotonergic neurons in the dorsal raphe nucleus scale O₂ utilization in response to traumatic stress": S5. Original WB

Original WB for Fig. 1

Fig. 1b-Human

NOX4

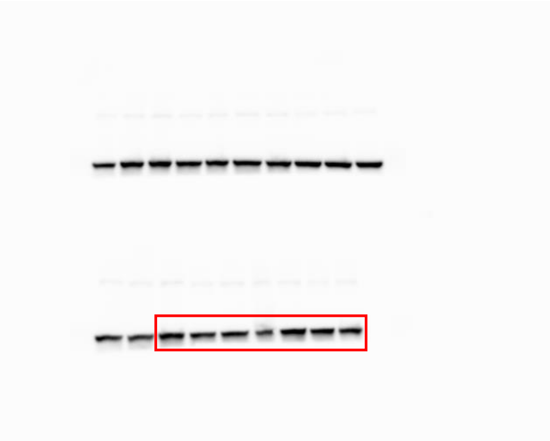

β-actin

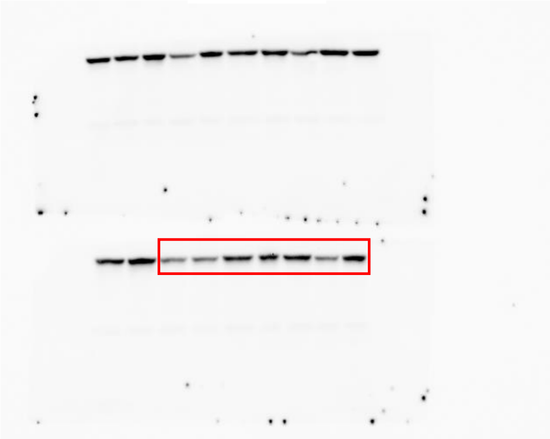

Fig. 1b-mice

NOX4 β-actin

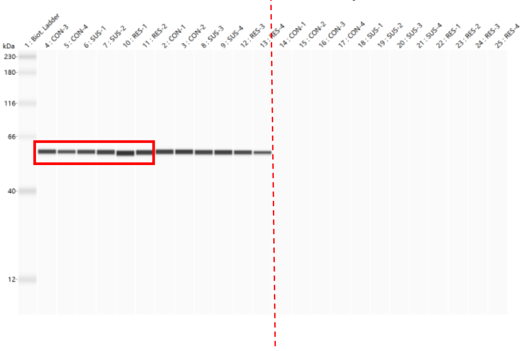

NOX4 β-actin

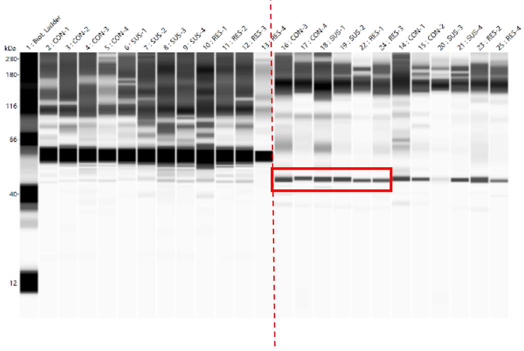

Original WB for Fig. 2

Fig. 2d

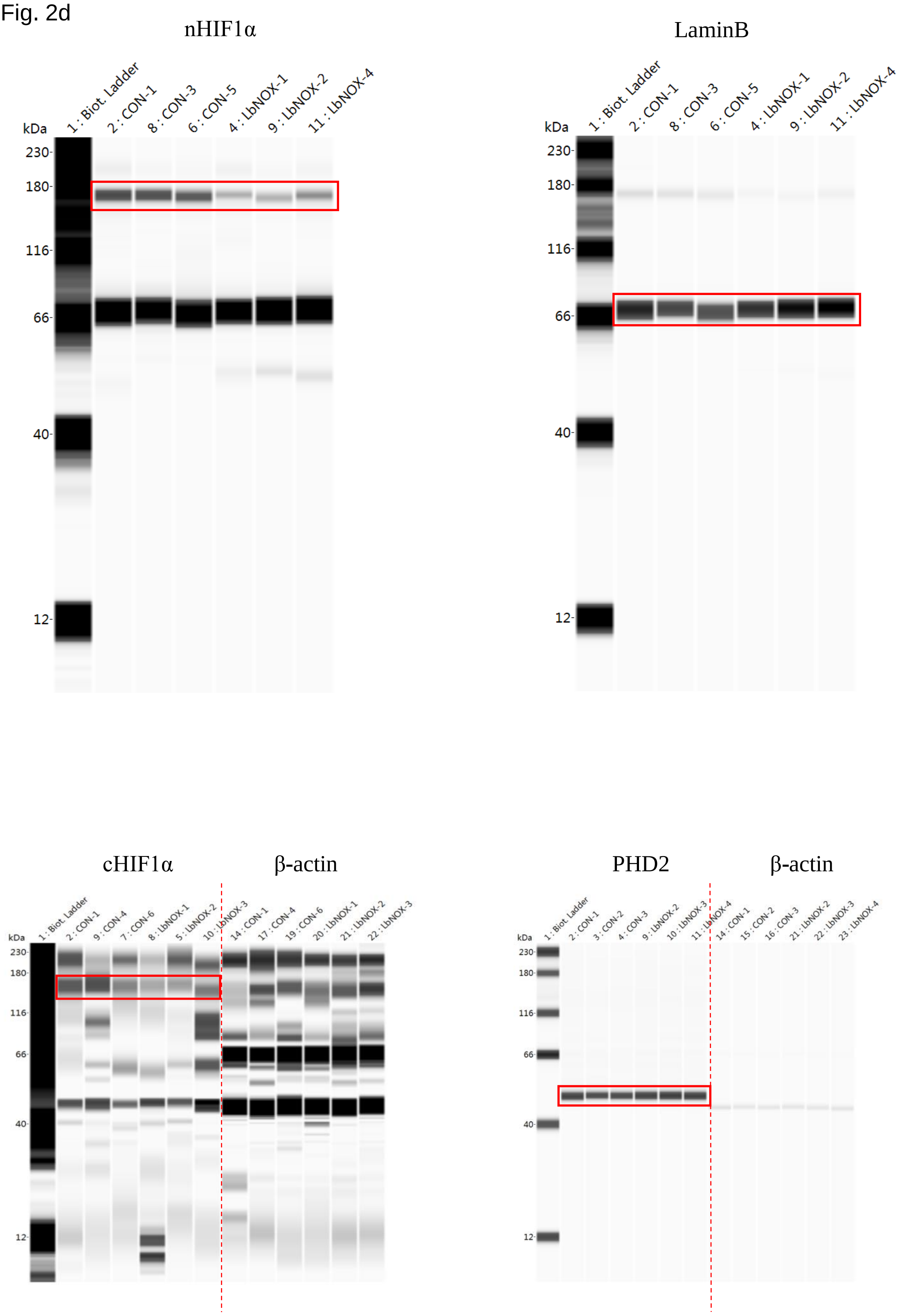

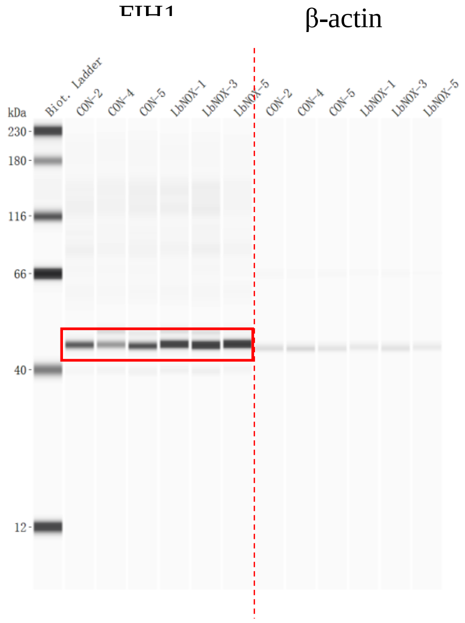
β-actin
kDa
Biot. Ladder
CON-2
CON-4
CON-5
LbNOX-1
LbNOX-3
LbNOX-5
CON-2
CON-4
CON-5
LbNOX-1
LbNOX-3
LbNOX-5
230
180
116
66
40
12

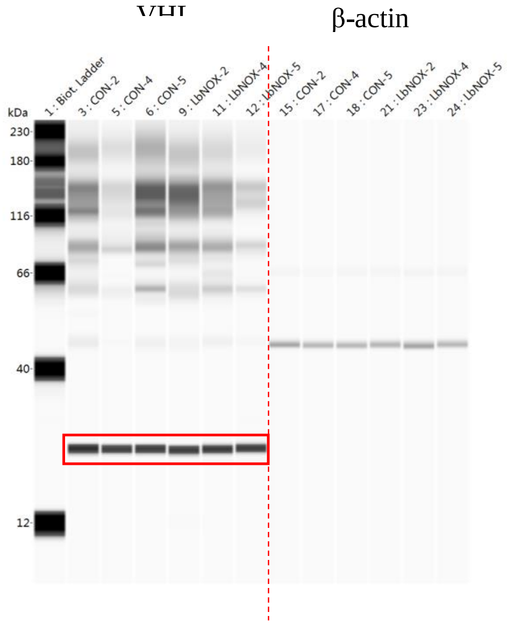
β-actin
kDa
1 : Biot. Ladder
3 : CON-2
5 : CON-4
6 : CON-5
9 : LbNOX-2
11 : LbNOX-4
12 : LbNOX-5
15 : CON-2
17 : CON-4
18 : CON-5
21 : LbNOX-2
23 : LbNOX-4
24 : LbNOX-5
230
180
116
66
40
12

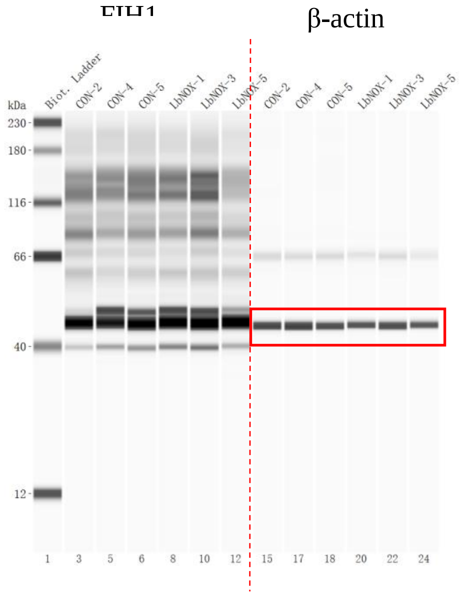
β-actin
kDa
Biot. Ladder
CON-2
CON-4
CON-5
LbNOX-1
LbNOX-3
LbNOX-5
CON-2
CON-4
CON-5
LbNOX-1
LbNOX-3
LbNOX-5
230
180
116
66
40
12
1 3 5 6 8 10 12 15 17 18 20 22 24

Original WB for Fig. 3

Fig. 3a for CSDS

nHIF1α

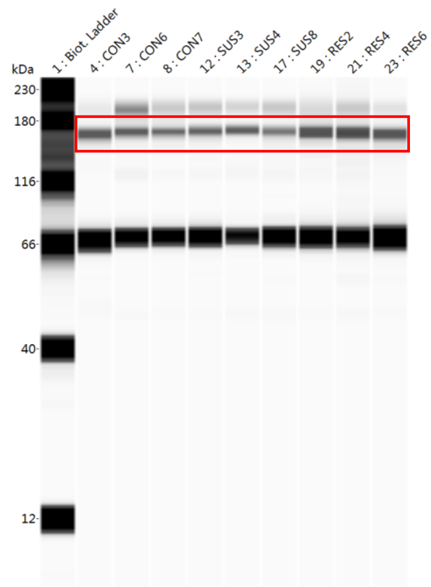


LaminB

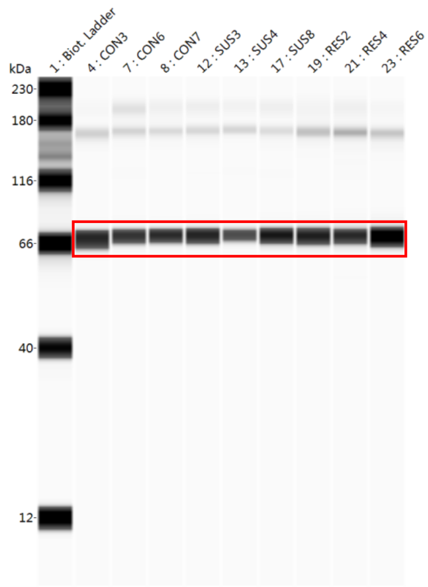


cHIF1α β-actin

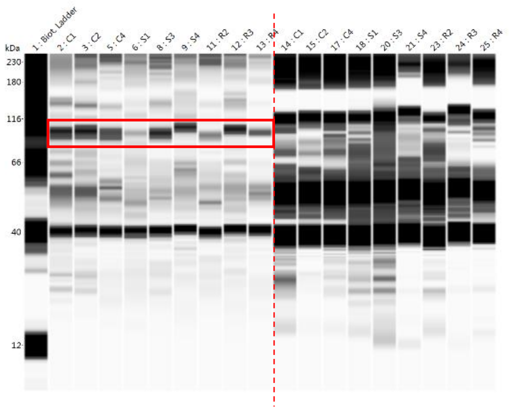


PHD2 β-actin

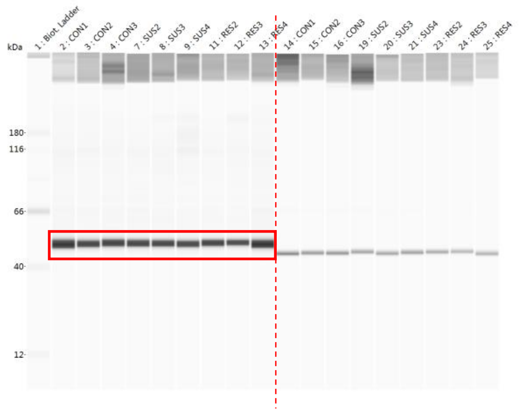


FIH1 β-actin

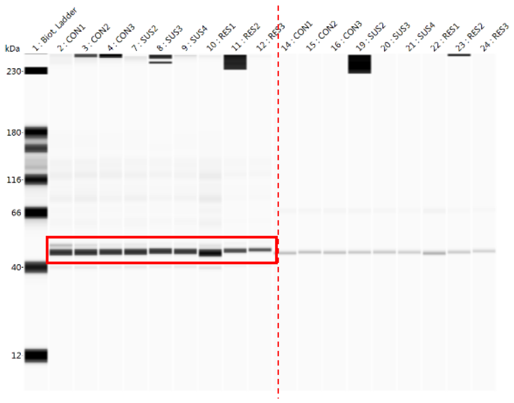


VHL β-actin

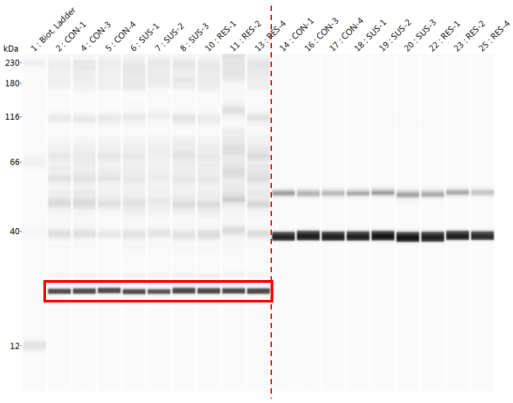

FIH1 β-actin

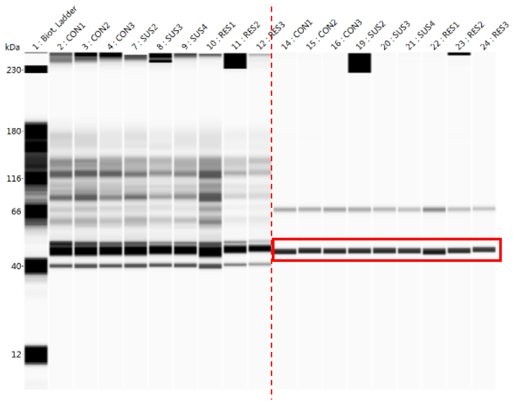
kDa
1 : Biot. Ladder
2 : CON1
3 : CON2
4 : CON3
7 : SUS2
8 : SUS3
9 : SUS4
10 : RES1
11 : RES2
12 : RES3
14 : CON1
15 : CON2
16 : CON3
19 : SUS2
20 : SUS3
21 : SUS4
22 : RES1
23 : RES2
24 : RES3
230
180
116
66
40
12

Fig. 3b for SEFL

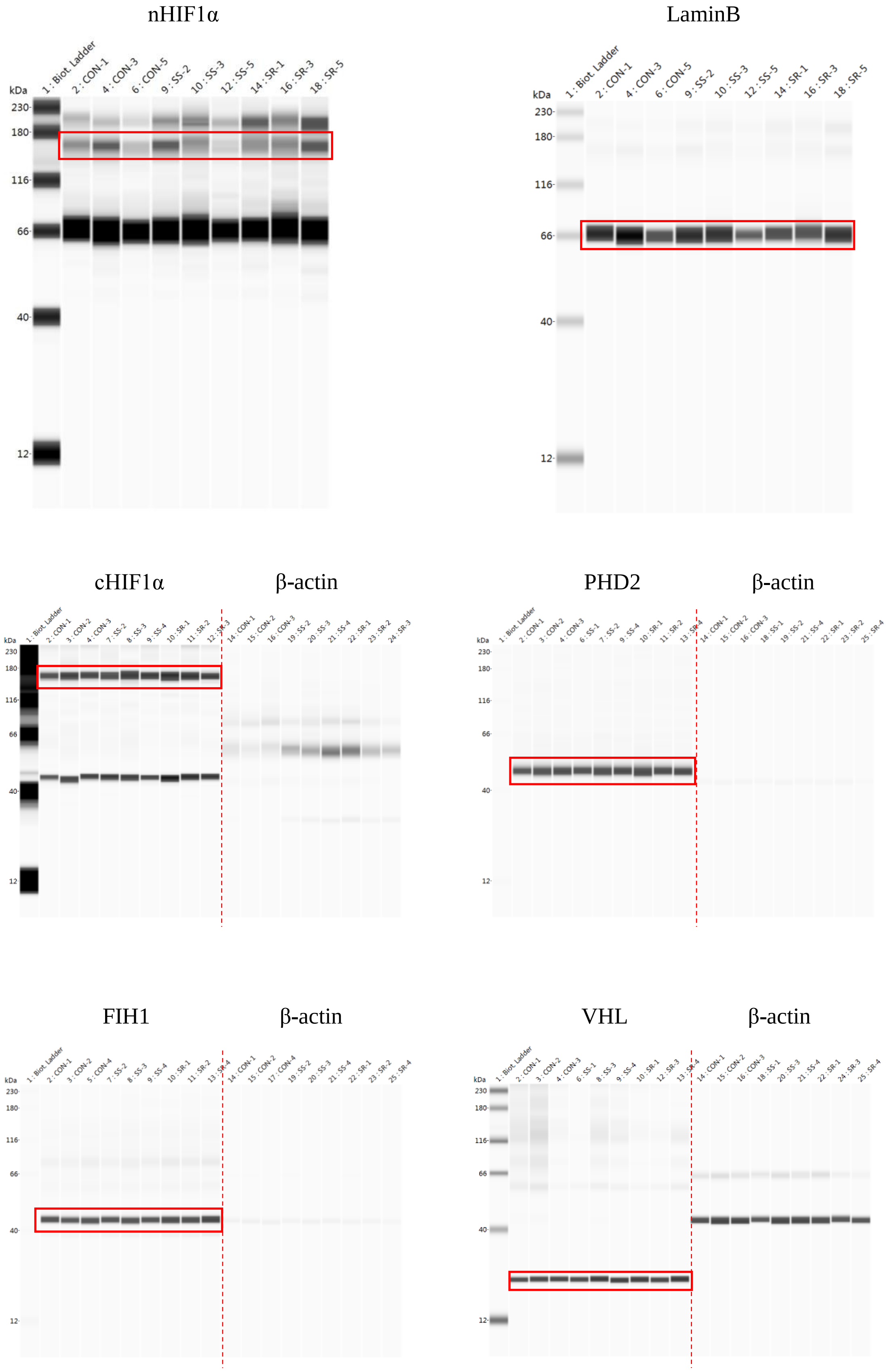

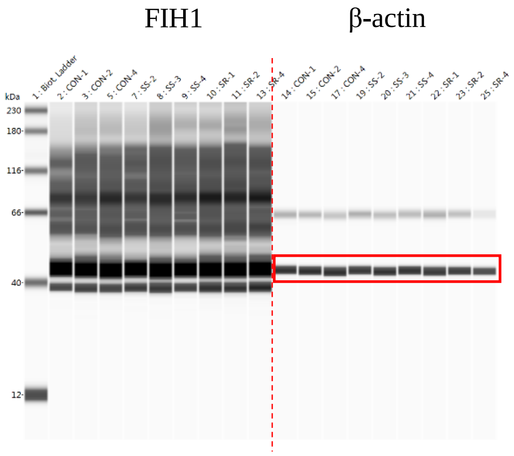
FIH1
β-actin
kDa
1 : Biot. Ladder
2 : CON-1
3 : CON-2
5 : CON-4
7 : SS-2
8 : SS-3
9 : SS-4
10 : SR-1
11 : SR-2
13 : SR-4
14 : CON-1
15 : CON-2
17 : CON-4
19 : SS-2
20 : SS-3
21 : SS-4
22 : SR-1
23 : SR-2
25 : SR-4
230
180
116
66
40
12

Fig. 3d

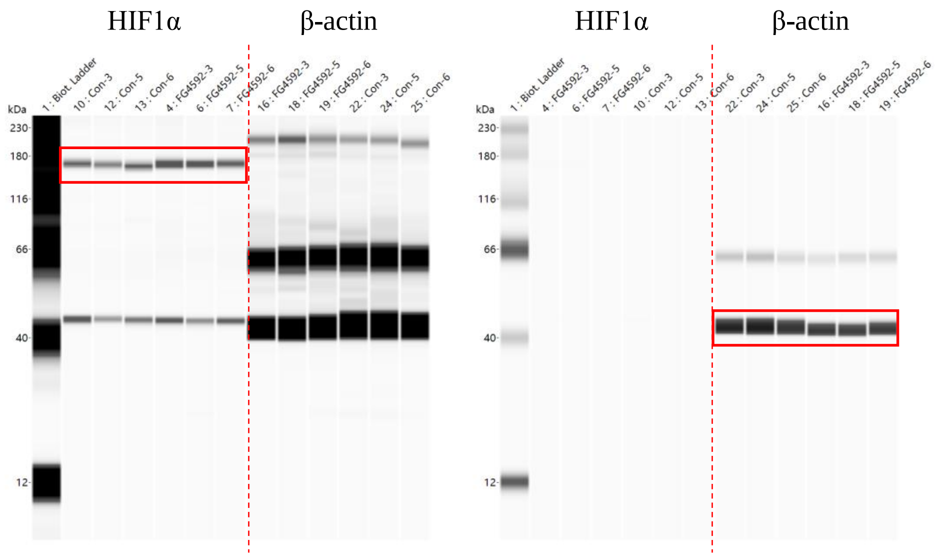
HIF1α
β-actin
1 : Biot. Ladder
10 : Con-3
12 : Con-5
13 : Con-6
4 : FG4592-3
6 : FG4592-5
7 : FG4592-6
16 : FG4592-3
18 : FG4592-5
19 : FG4592-6
22 : Con-3
24 : Con-5
25 : Con-6
kDa
230
180
116
66
40
12
HIF1α
β-actin
1 : Biot. Ladder
4 : FG4592-3
6 : FG4592-5
7 : FG4592-6
10 : Con-3
12 : Con-5
13 : Con-6
22 : Con-3
24 : Con-5
25 : Con-6
16 : FG4592-3
18 : FG4592-5
19 : FG4592-6
kDa
230
180
116
66
40
12

Original WB for Fig. 7

Fig. 7a

NOX4 GPX4

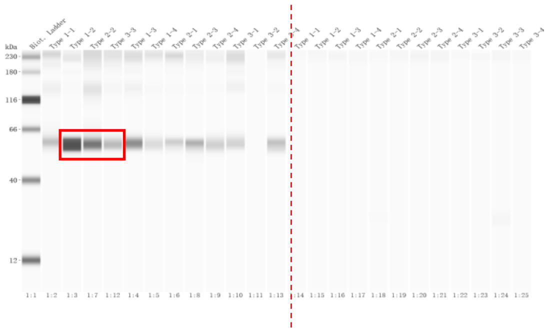

NOX4 GPX4

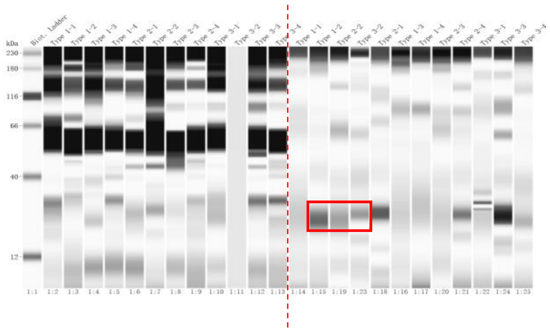

GAPDH

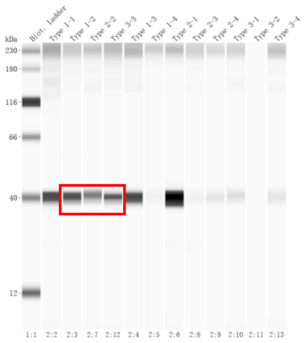

### Fig. 7d for Type I neurons

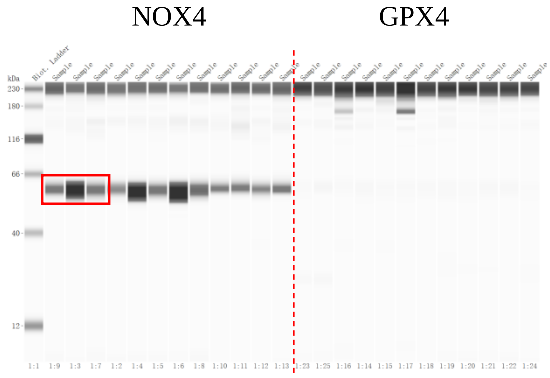


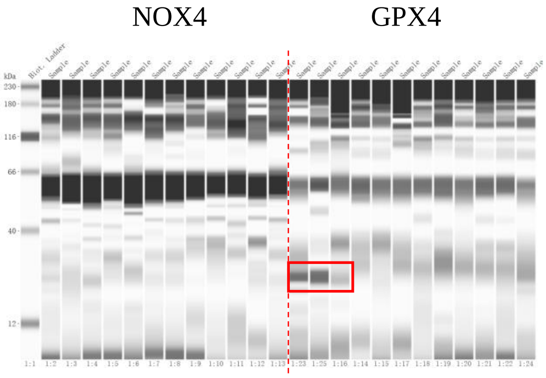


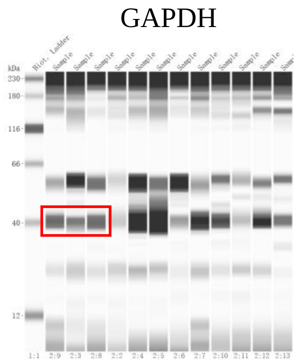


### Fig. 7d for Type II neurons

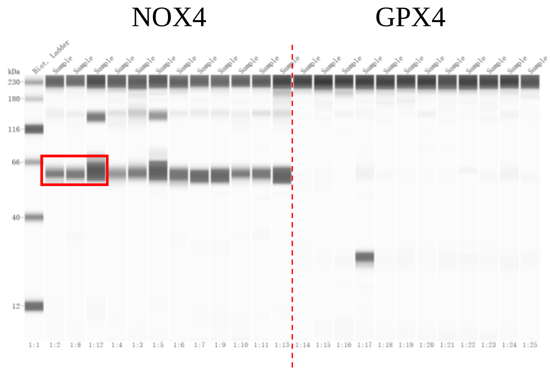


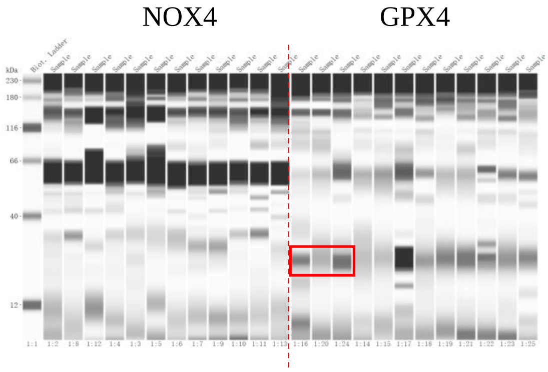


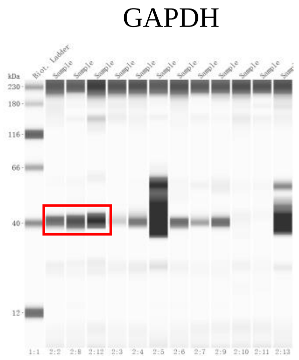

Fig. 7d for Type III neuron

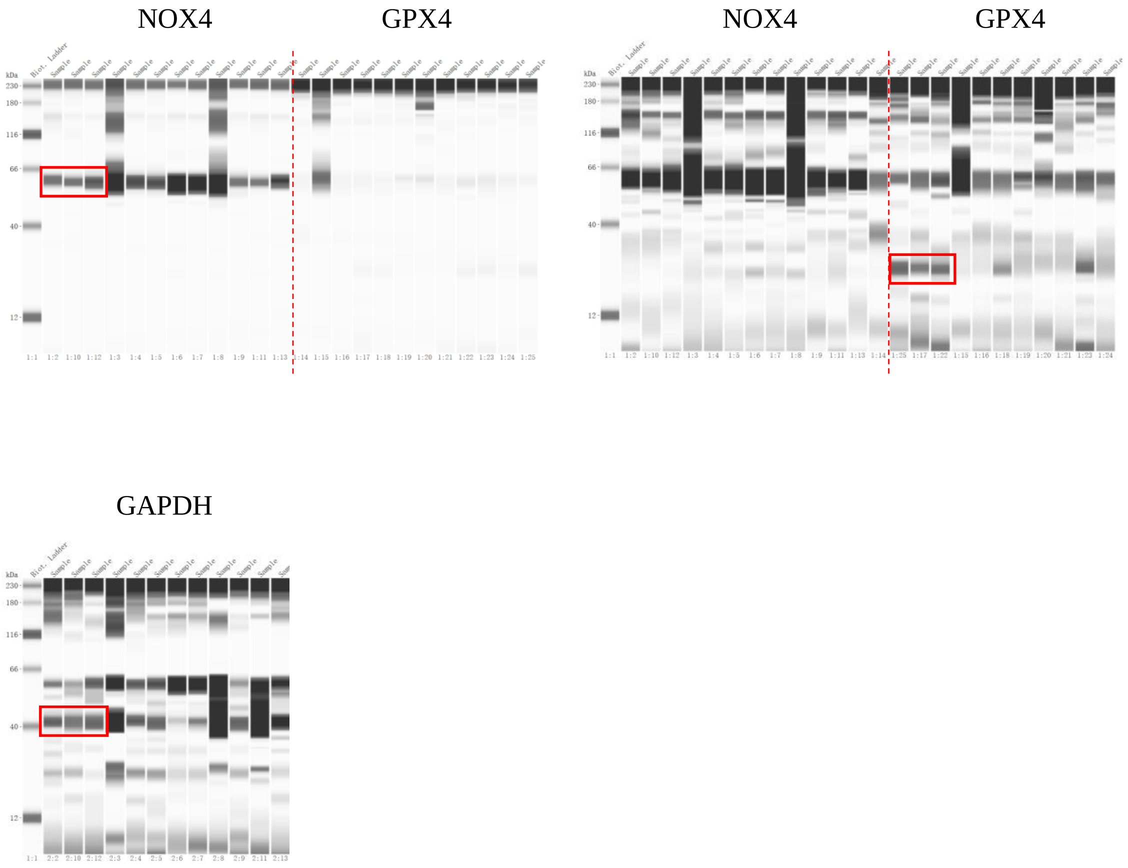

Original WB for Fig.8

Fig. 8b

NOX4 β-actin

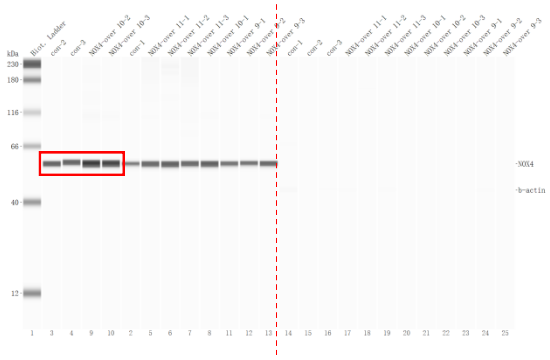

NOX4 β-actin

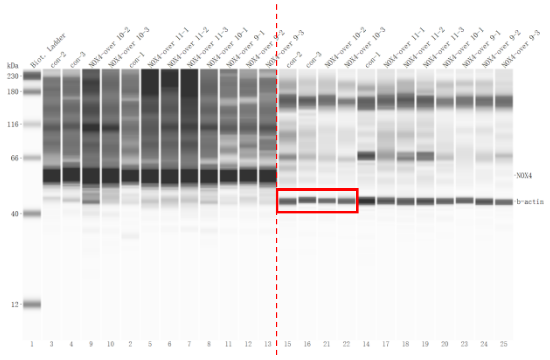

Original WB for Supplementary Fig. 4

Supplementary Fig. 4b for *Hif1an*-shRNA injection

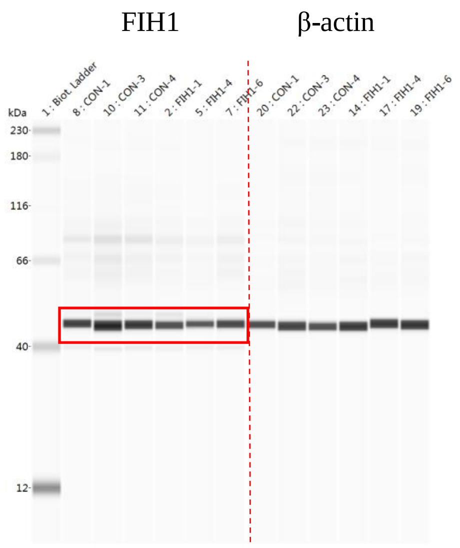


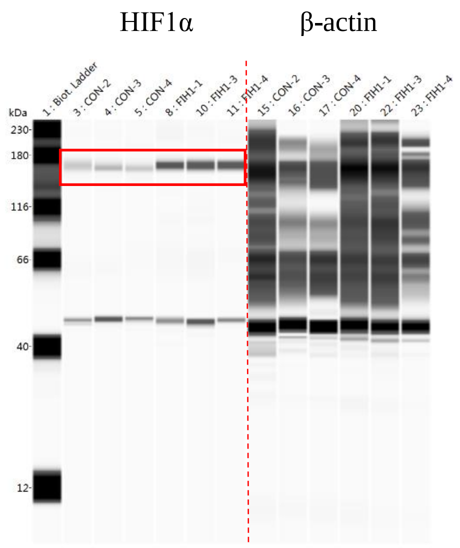


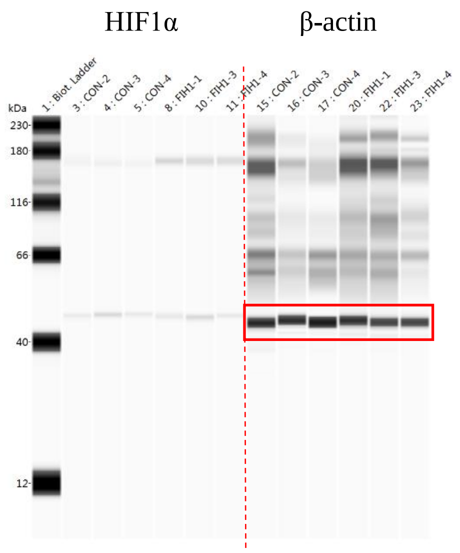

Supplementary Fig. 4b for *Egln1*-shRNA injection

PHD2 β-actin

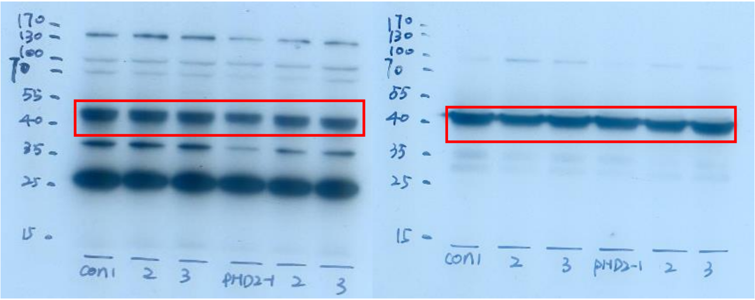

HIF1α β-actin HIF1α β-actin

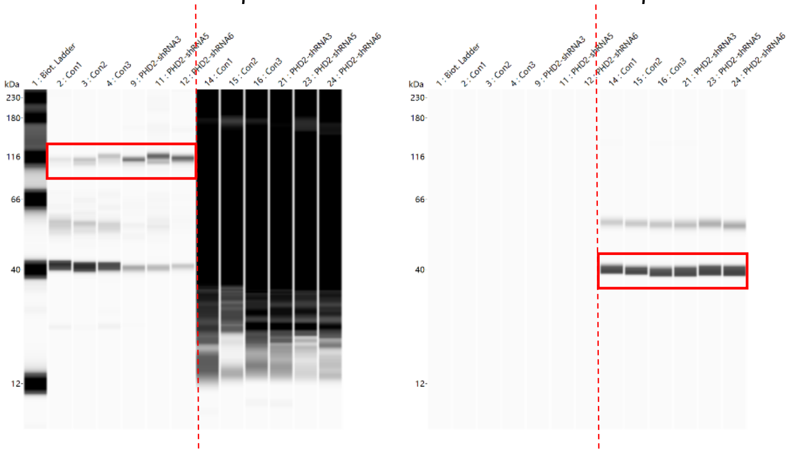

### Original WB for Supplementary Fig. 10

#### Supplementary Fig. 10d

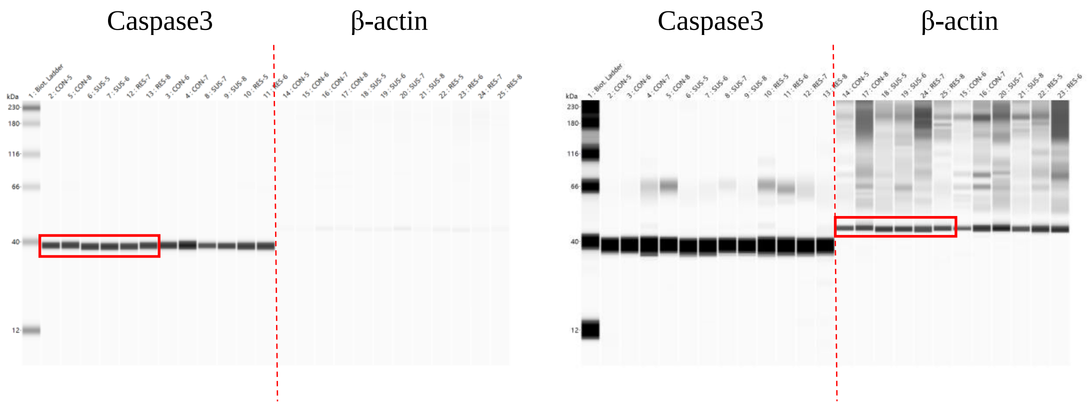
